# Single cell transcriptomics identifies master regulators of neurodegeneration in SOD1 ALS motor neurons

**DOI:** 10.1101/593129

**Authors:** Seema C. Namboori, Patricia Thomas, Ryan Ames, Sophie Hawkins, Lawrence O. Garrett, Craig R. G. Willis, Alessandro Rosa, Lawrence W. Stanton, Akshay Bhinge

## Abstract

**Background:** Bulk RNA-Seq has been extensively utilized to investigate the molecular changes accompanying motor neuron degeneration in Amyotrophic Lateral Sclerosis (ALS). However, due to the heterogeneity and degenerating phenotype of the neurons, it has proved difficult to assign specific changes to neuronal subtypes and identify which factors drive these changes. Consequently, we have utilized single cell transcriptomics of degenerating motor neurons derived from ALS patients to uncover key transcriptional drivers of dysfunctional pathways.

**Results:** Single cell analysis of spinal neuronal cultures derived from SOD1 E100G ALS and isogenic iPSCs allowed us to classify cells into neural subtypes including motor neurons and interneurons. Differential expression analysis between disease and control motor neurons revealed downregulation of genes involved in synaptic structure, neuronal cytoskeleton, mitochondrial function and autophagy. Interestingly, interneurons did not show similar suppression of these homeostatic functions. Single cell expression data enabled us to derive a context-specific transcriptional network relevant to ALS neurons. Master regulator analysis based on this network identified core transcriptional factors driving the ALS MN gene dysregulation. Specifically, we identified activation of SMAD2, a downstream mediator of the TGF-β signalling pathway as a potential driving factor of ALS motor neuron degeneration. Our phenotypic analysis further confirmed that an activated TGF-β signalling is major driver of motor neuron loss in SOD1 ALS. Importantly, expression analysis of TGFβ target genes and computational analysis of publicly available datasets indicates that activation of TGFβ signalling may be a common mechanism shared between SOD1, FUS and sporadic ALS.

**Conclusions:** Our results demonstrate the utility of single cell transcriptomics in mapping disease-relevant gene regulatory networks driving neurodegeneration in ALS motor neurons. We find that ALS-associated mutant SOD1 targets transcriptional networks that perturb motor neuron homeostasis.

## Introduction

Amyotrophic Lateral Sclerosis (ALS), also known as Lou Gehrig’s disease, is an age-onset fatal neurodegenerative disorder that affects motor neurons in the brain and spinal cord[1]. Patients display progressive paralysis and eventually die due to failure of the respiratory muscles, commonly within 3-5 years of diagnosis[2]. Despite extensive research, the causes underlying the observed degeneration are incompletely understood. Hence, so far, there is no cure for ALS. Understanding the molecular drivers of neurodegeneration in ALS can potentially lead to the development of life-saving therapies. Approximately 20% of ALS cases are familial with mutations identified in genes spanning diverse cellular functions including SOD1[3]. Animal models have been extensively used to study ALS and have revealed important insights into disease mechanisms[4]. However, species-specific differences and the need to overexpress the mutant proteins to generate phenotypes have necessitated developing human models of the disease that can complement existing animal models[5]. Induced pluripotent stem cells (iPSC) derived from patients suffering with ALS provide a powerful model to study ALS. ALS patient-derived iPSC bear the disease-causing mutations in a physiologically relevant background and can be readily differentiated into clinically relevant neurons[6]. Such diseased neurons can now be compared to healthy neurons to model key aspects of the disease such as neuron survival, morphometric defects, electrophysiological dysfunctions and protein/RNA aggregation foci *in vitro*[7–12].

Molecular characterisation of these neurons using “omics” tools has uncovered important insights into disease pathophysiology[8, 9, 13, 14]. However, application of genomic tools such as RNA-sequencing to ALS neurons in bulk has serious drawbacks. Current differentiation protocols generate motor neurons at efficiencies ranging from 50-80% with the efficiency varying depending upon the iPSC line used. The differentiated neurons are usually a mix of motor neurons (MN) and “non-motor neurons”, typically spinal interneurons (IN). Additionally, long term cultures of these neurons that are required for phenotypic characterization commonly display some degree of glial cell proliferation. Importantly, ALS motor neurons display progressive degeneration. This suggests that at any given time point, neurons can be expected to be in different stages of degeneration and hence may display a differential expression of key drivers of disease pathology. Bulk RNA analysis of these cultures would not only average cell type expression but also the motor neuron specific degeneration expression signatures. To address these issues, we performed single cell transcriptomic analysis of degenerating ALS SOD1 mutant neurons. Our analysis reveals MN specific transcriptional networks regulating synaptic function and neuronal cytoskeleton to be downregulated in ALS MN. Importantly, single cell data enabled us to build a disease relevant transcriptional network that was used to identify key transcription factors driving the ALS-associated gene dysregulation.

## Materials and Methods

### Human iPSC culture

ALS patient-derived iPSC bearing SOD1 E100G/+ (ND35662) mutation were obtained from the Coriell Institute for Medical Research. ALS iPSC and the genome edited isogenic control iPSC were maintained as colonies on human ES qualified matrigel (Corning) in mTeSR (StemCell Technologies). Colonies were routinely passaged in a 1:6 split using Dispase. Mycoplasma testing was conducted regularly to rule out mycoplasma contamination of cultures.

### Differentiation of iPSC into spinal motor neurons

iPSC were plated as colonies onto matrigel and differentiated by treatment with neuronal differentiation media (DMEM/F12:Neurobasal in a 1:1 ratio, HEPES 10mM, N2 supplement 1%, B27 supplement 1%, L-glutamine 1%, ascorbic acid 5uM, insulin 20ug/ml) supplemented with SB431542 (40uM), CHIR9921 (3uM) and LDN8312 (0.2uM) from day 0 till day 4. Cells were caudalized by treatment with 0.1uM retinoic acid starting at day 2 and ventralized with 1uM purmorphamine starting at day 4 and continued till day 10. At day 10, OLIG2 positive motor neuron progenitors were re-plated onto poly-D-lysine/laminin coated wells and differentiated by treating the cells with N2B27 media supplemented with BDNF 20ug/ml, GDNF 10ug/ml and DAPT 10uM. DAPT treatment was stopped at day 14 and neuronal cultures were pulsed with mitomycin at a dose of 10ug/ml for 1 hour to prevent further proliferation of any undifferentiated progenitors. Neuronal cultures were maintained by changing media every other day.

### MN survival assay

MN were differentiated from SOD1 E100G and the isogenic control iPSC as described above in 96-well optically clear tissue culture plates. Day 30 cultures were fixed and stained for ISL1 to assess MN counts. A separate plate cultured under the same condition was allowed to proceed till day 44, when it was fixed and stained for ISL1. Nuclei were stained using Hoechst 33342. MN counts at day 40 were compared with day 30 to assess MN survival. We performed three independent differentiations with two technical wells used per replicate. Data from the technical wells was pooled to generate counts for each replicate.

### Neuronal survival assays in response to TGFβ perturbation

We found that the process of immunostaining occasionally resulted in neuronal detachment leading to underestimates of neuronal counts. Hence, cultures that showed neuronal detachment were discarded. We realized that a better approach was to treat live neurons with the cell-permeable nuclear dye Hoechst 33342 and image the same well at different time points. Given that almost 80% of the cells in culture were motor neurons, nuclei counts were expected to closely approximate MN counts. Neuronal cultures at day 30 were treated with Hoechst 33542 (0.25 ug/ml) for 45 minutes and media was replaced with standard neuronal culture media as described above. Nuclear stained cultures were analysed by live imaging in the DAPI channel at day 30 and again at day 40. This allowed us to assess nuclear counts for the same well at the two different time points. Day 40 counts were compared with day 30 counts to assess percentage loss of neurons using 3 independent differentiations with two wells as technical replicates per treatment per differentiation. SB431542 was dissolved in DMSO. Hence, a DMSO only treatment was used as control for the TGFβ inhibiton experiments. The final concentration of DMSO was maintained at 0.1%. TGFβ1 was resuspended in PBS and we used neurons treated with PBS as control. Apoptosis was estimated in independent neuronal cultures using the Promega RealTime Glo apoptosis assay according to the manufacturer’s instructions.

### Immunofluorescence

Cells were fixed with 4% paraformaldehyde, permeabilized with ice-cold methanol for 5 minutes and washed with PBS containing 10% serum for 1 hour at room temperature. Cells were incubated with primary antibodies (Table S1) diluted into PBS containing 10% serum and incubated overnight at 4°C. Next day, cells were washed and incubated with Alexa-fluor conjugated secondary antibodies (Molecular probes) for 45 minutes at room temperature and nuclei were stained with Hoechst 33542 (Molecular probes). Images were obtained in an automated fashion on the ImageXpress Pico (Molecular devices). Nuclear identification and p-SMAD2 nuclear signal was quantified in an automated fashion using the imageXpress software. Threshold intensities were maintained the same across wells.

### Quantitative RT-PCR

Total RNA was extracted with the miRNeasy kit (Qiagen) and reverse transcribed using random hexamers and the High Capacity reverse transcription system from Applied Biosystems. Quantitative PCR was performed using the SYBR GREEN PCR Master Mix from Applied Biosystems. The target gene mRNA expression was normalized to the expression of two housekeeping genes (HPRT1 and RPL13), and relative mRNA fold changes were calculated by the ΔΔCt method. Primer sequences are included in Table S2.

### Single-cell capture and library preparation

Single cells were captured using standard protocol of C1 single-cell auto prep system (Fluidigm). Two independent differentiations were set up a few days apart. At day 44, differentiated neuronal cultures were dissociated into single cells by Accutase and loaded onto the C1 chip. We used one chip per genotype per differentiation. Chips for each replicate (one for the SOD1 E100G and the other for the isogenic control) were loaded in parallel into two separate machines. We used a total of four chips labelled A, B, C and D. Chips A and B captured the isogenic control MN while chips C and D captured the SOD1 E100G MN. Post cell capture, each well of the chip was manually inspected to identify wells bearing single cells. Next, lysis, reverse transcription and PCR amplification of the cDNA was perform in an automated fashion within the C1 instrument. To prepare single-cell libraries, cDNA products from each single cell were harvested from C1 chip followed by concentration and quantification using PicoGreen dsDNA Assay kit. Sequencing libraries were generated using Illumina Nextera XT library preparation kit.

### Read processing, mapping and quality control

Fastq files were processed using Salmon with a partial decoy index for human gene annotation GENCODE release 19 [15]. Transcript level counts were collapsed to generate read counts per gene using the tximport R package. This yielded 33,000 transcripts across 365 libraries. Only libraries deemed to be single cells were retained for further analysis (332 cells). Additionally, libraries were filtered for low quality cells using PCA as described in the main text. A gene was deemed to be poorly expressed if it was found to be present in less than 10 cells at a read count threshold of 2. The filtering process yielded 332 single cells and 14774 genes for further analysis.

Single cell dissociation and harvest protocols lead to lysis of cells that release their cellular mRNA into the mixture. This RNA termed “ambient mRNA” is released from dying cells and has been observed in the absence of cell capture in droplet based platforms[16]. The ambient mRNAs typically arise from genes expressed at high levels. We observed that some of the microfluidic chambers that were marked as empty had generated RNA reads that mapped to human transcripts. It is possible that these chambers contained a cell that was missed by the human observer. But we decided to treat transcript counts arising from empty chambers as ambient mRNA. For each plate, we summed up the counts per gene across the empty chambers and estimated counts per million (cpm) reads for each gene. Next, we averaged the cpm values per gene across the two replicates plates for the healthy and ALS samples separately. We marked all genes at a cpm threshold above 100 as being affected by ambient RNA expression. We performed this filtering for the healthy and ALS samples individually to avoid excluding genes that were lowly expressed in either the healthy or the ALS samples. This resulted in a total of 1120 genes being marked as affected by the ambient mRNA. The ambient geneset was excluded from the clustering analysis to identify neural subtypes. Additionally, we excluded these genes from the sorted differentially expressed geneset used in the master regulator analysis (see below). This genelist has been included as Supplementary data file S1.

### Differential gene expression analysis

We used DESeq2 with parameters optimized for single cell data analysis using the command: DESeq(dds, test = “Wald”, sfType = “poscounts”, minReplicatesForReplace = Inf, useT = T, minmu = 1e-6). The design used for the analysis was Batch + Sample, where Batch indicated the replicate and Sample indicated the genotype.

### kNN smoothing of single cell read counts

We adapted the algorithm described by Wagner et al[17]. Briefly, the read count smoothing algorithm worked as follows: 1) We started with a matrix of N cells X m genes where N=189 and m=14774 2) Count data was variance stabilized by taking the logarithm of the counts after adding 1 for each value in the single cell matrix. This was because, for our dataset, the log transformation worked better than the Freeman Tukey transform used in the original study. 3) Log transformed read counts were quantile normalized. 4) Euclidean distances between the cells were calculated using the first 8 principal components. The threshold of 8 is determined empirically using a scree plot. 4) For each cell, we calculated a weighted average of the read counts per gene between that cell and its k neighbours. The weights for the averaging are assigned as the inverse square root of the Euclidean distance between the cell and its neighbour. This ensured that neighbours that were far away in Euclidean space did not contribute as much to the final smoothened count than the nearest neigbours. The value of k is iteratively increased starting from 1 according to the following equation: k = min(2^step^ – 1, kmax); kmax = sqrt(N); step = 1:max_steps; max_steps = floor(log2(kmax + 1)); For example, in our study, for N=189, we get kmax=14 and max_steps=3.0. This results in 3 iterations where the value if k equals 1, 3 and 7 in each iteration successively. The final output is a matrix of counts that are quantile normalized and log transformed.

### Weighted gene coexpression network analysis

To identify functional modules of genes associated with ALS we used weighted gene coexpression network analysis (WGCNA) implemented in the R statistical language. K-NN smoothened normalized read counts were used to build a coexpression network. The coexpression network was constructed with WGCNA using a soft thresholding power of 6 using the signed-hybrid approach. Modules in the network were identified using the *cutTreeDynamic* function with a minimum module size of 50. Modules with similar expression profiles were merged if their eigengene correlation coefficients were >=0.75. We used Pearson’s correlation coefficient to assess associations between module eigengenes and disease state or neuronal subtypes. *P-values* were corrected using the method of Benjamini and Hochberg and correlations with an adjusted p-value < 0.01 were deemed significant. Gene ontology enrichment analysis of disease associated modules was carried out using the anRichment R package.

### Master regulator analysis

The master regulator analysis was performed using the RTN R package. Smoothened counts for 189 neurons were used as input to build a transcriptional network for 1137 TFs present in the dataset. TF annotations were obtained from AnimalTFDB. P-values for network edges were estimated from a pooled null distribution using 1000 permutations. Edges with a p-value < 5e-8 were retained for MRA. The predicted targets of each TF were termed as the regulon. The regulon for each TF was classed as positive or negative based on the Pearson correlations. To identify master regulators, the differential gene expression between ALS and control MN (after removing the ambient geneset) was used as a phenotype and sorted from most upregulated to most downregulated. The RTN package was used to conduct a GSEA like analysis to identify whether a TF regulon (positive or negative) was enriched towards one end of the sorted list of differentially expressed genes. P-values were estimated based on 1000 permutations of the dataset and adjusted using the Benjamini Hochberg method. Gene ontology enrichment analysis of the regulons was carried out using the anRichment R package.

### Analysis of publicly available ALS datasets

Normalized read count data for GSE54409 (human SOD1 A4V iPSC derived MN purified using flow sorting based on the HB9 reporter)[8] were downloaded from the gene expression omnibus(GEO). P-values were estimated by performing a t.test per gene. Read counts were averaged across replicates per gene and log2 transformed. Fold changes were estimated by subtracting the log2 counts for the ALS and isogenic controls: log2(ALS) – log2(control). Microarray expression values for GSE46298 (laser-capture microdissected MN from spinal tissue obtained from the mouse SOD1 G93A ALS model)[18] and GSE18920 (laser-capture microdissected MN from spinal tissue obtained from sporadic ALS patients post-mortem)[19] were downloaded from the gene expression omnibus. Expression values were background subtracted, normalized and log transformed using RMA from the affy R package[20]. Only genes with median expression values above background were included in the analysis. Probes mapping to the same gene were collapsed. Differential expression analysis was performed using the limma package in R[21]. Differentially expressed genes for GSE76220 (laser-capture microdissected MN from spinal tissue obtained from sporadic ALS patients post-mortem) [13] were obtained from the supplementary data for that study. This study had removed genes activated by the wound healing response by excluding genes with the highest principal component (PC1) eigengene values. We obtained both, the full (all genes) and the filtered (PC1 adjusted genes) datasets. Count data for GSE140747[22] (motor neuron development gene expression: D0-D15) and GSE98288[23, 24] (D21, D35 vs iPSC) was downloaded from the gene expression omnibus. For GSE98288, we only used counts from the control iPSC differentiations. P-values for all datasets were corrected for multiple hypotheses using the Benjamini Hochberg procedure. Differentially expressed genes identified from our bulk RNA-seq analysis of SOD1 E100G iPSC derived MN were obtained as described previously[25]. To sort genes, each gene was assigned a score = −1*log10(p-value) * sign(fold change). Genes were sorted based on this score in a decreasing manner so that the most significantly up regulated genes were assigned to the top of the list.

## Results

### Single cell RNA-Seq analysis of ALS and control neurons

We previously developed an *in vitro* model of ALS MN degeneration where MN differentiated from mutant SOD1 iPSC display disease-specific phenotypes such as reduced cell survival and morphometric defects compared to their isogenic control counterparts[25]. We differentiated iPSC derived from patients bearing the SOD1 E100G mutation, as well as the corresponding CRISPR edited isogenic control (SOD1 E100E) into MN (Fig.1a) [25]. Mature MN appeared by day 30 and expressed MN markers such as ISL1 and NF-H in addition to the pan-neuronal marker MAP2 (Fig. 1b). Using our protocol, both ALS and control iPSC could be efficiently and reproducibly differentiated into MN at similar efficiencies (75-77% ISL1+ cells across five independent differentiations) (Fig. 1b). To assess for a survival phenotype, we differentiated the ALS SOD1 and isogenic control iPSC into MN, and assayed for MN survival between day 30 and day 44. ALS iPSC derived neurons displayed a 40% loss in survival compared to the isogenic control neurons similar to our previous observations[25] (Fig. 1c). To gain deeper insights into the mechanisms driving neurodegeneration, we performed single cell RNA-sequencing on the ALS and isogenic control MN at day 44 of our differentiation protocol (Fig. 1a). Neuronal cultures from two independent differentiations were dissociated into single cells, captured and lysed in a fully automated fashion using a microfluidic platform followed by library preparation and deep sequencing of transcripts in the individual cells (Methods). We captured >80 single cells per genotype per replicate. Overall, we captured a total of 332 single cells that included 165 cells from the ALS cultures and 167 cells from the isogenic controls.

**Fig. 1.**
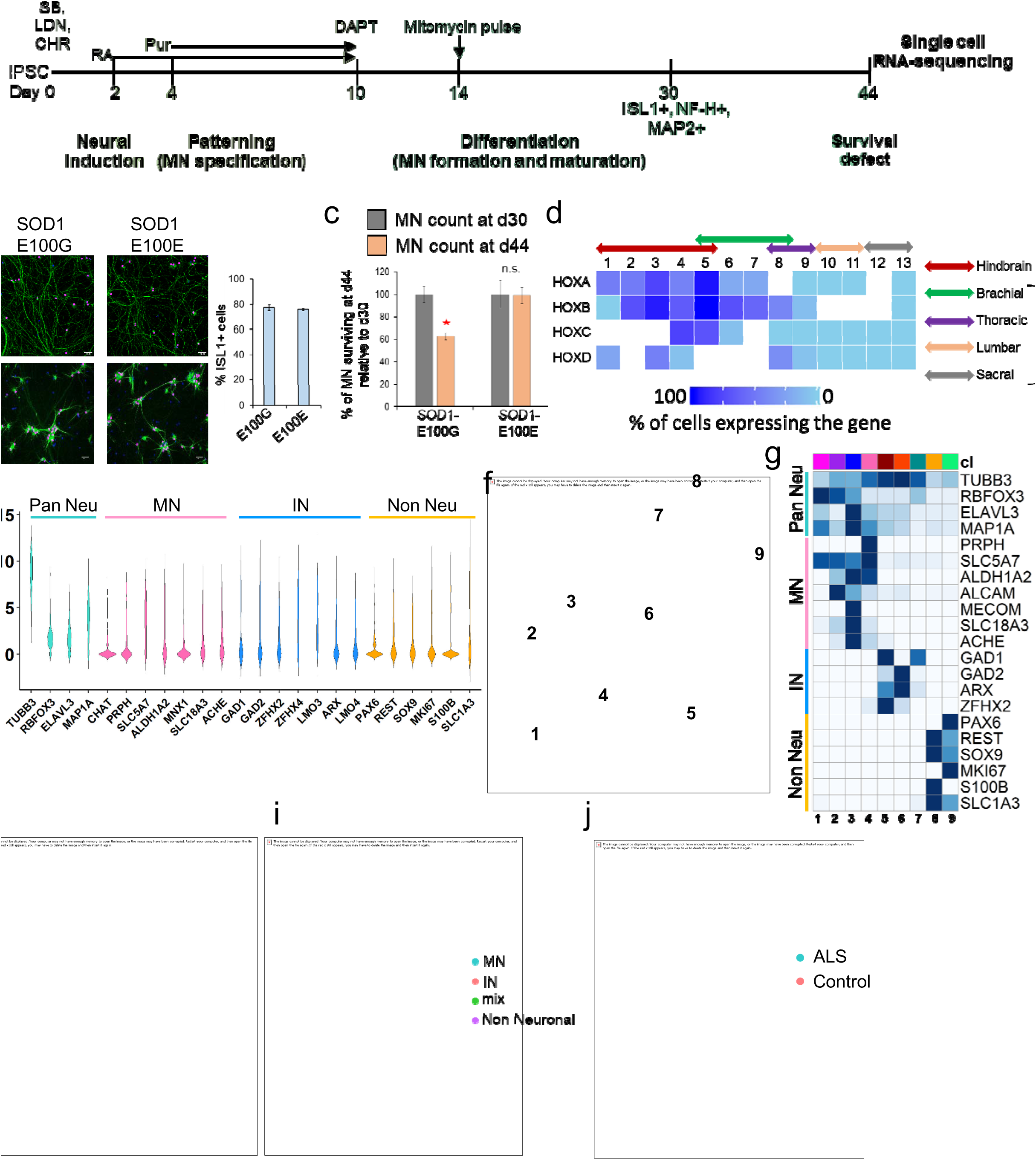
Single cell RNA-sequencing of iPSC-derived neurons. a) Schematic depicting the differentiation protocol used to derive MN from human iPSC. The horizontal line depicts the timeline of differentiation with the numbers indicating days. Day 0 indicates the first media change to induce differentiation. SB: SB431542, LDN: LDN193189, CHR: CHIR99021, RA: Retinoic acid, Pur: Purmorphamine. b) iPSC-derived MN at day 30 stain positive for the MN markers ISL1 and NF-H as well as the pan-neuronal marker MAP2. Scale bar indicates 50µm. c) MN (ISL1+) were counted at d30 and d44 of the differentiation protocol. D44 MN counts were normalized to d30 counts. SOD1 E100G indicates the MN derived from mutant SOD1 iPSC. SOD1 E100E indicates the isogenic corrected control MN. Error bars shown are s.e.m. n=3 independent biological replicates. * indicates p-value < 0.01. n.s. indicates not significant. d) Heatmap displaying % of cells expressing specific HOX genes. Read count threshold was set at 2. White space indicates that the corresponding HOX paralog is not expressed in humans. Coloured solid arrows indicate the HOX code for specific spinal segments along the rostro-caudal axis. e) Violin plots displaying distribution of expression levels of neuronal markers (TUBB3, RBFOX3, ELAVL3, MAP1A), MN markers (CHAT, PRPH, SLC5A7, ALDH1A2, MNX1, SLC18A3 or VaCHT, ACHE), interneuron markers (GAD1, GAD2, ZFHX2, ZFHX4, LMO3, ARX, LMO4) and non-neuronal markers (PAX6, REST, SOX9, MKI67, S100B, SLC1A3) across all cells. f) Uniform manifold approximation and projection (UMAP) plot showing clustering of single cells into nine clusters. g) Normalized mean expression of neural markers across all nine clusters (cl). Clusters have been coded by numbers (at the bottom) and by colour (at the top). These correspond to the numbers and colours shown in panel f. Clusters 1-4 express high levels of MN markers, clusters 5-7 express IN markers, while clusters 8,9 express glial and progenitor markers. h) Partition analysis shows the MN cluster 4 associating closely with IN clusters 5,6 while IN cluster 7 associating with non-neuronal clusters 8,9. Hence, clusters 4 and 7 were deemed to be mixed clusters i.e. cells that cannot be classified unambiguously as IN or non-neuronal cells. i) UMAP plot showing classification of single cells into MN, IN and non-neuronal cells based on marker expression. j) UMAP plot showing distribution of the identified cell types amongst the ALS SOD1 E100G (ALS) and isogenic SOD1 E100E control (Control) samples.

Single cell transcriptomes were sequenced to an average depth of 1.5e6 reads per cell and the overall sequencing depth was similar across the ALS and control datasets (Fig. S1a). The proportion of total reads mapped to the genome is an indicator of library quality[26]. Both ALS and control cells displayed similar mapping proportions indicating there was no bias towards poor quality cells in any one dataset (Fig. S1b). At a read count threshold of 1.0, both datasets expressed an average of ∼8000 genes (Fig. S1c). To remove poorly amplified RNA libraries (indicated by low mapping ratios and low numbers of expressed genes) and lowly expressed genes, single cell libraries were subjected to a set of quality control criteria that included: 1) total mapped reads, 2) percentage mapped reads, 3) percentage of mitochondrial reads, 4) number of genes expressed (Methods). To identify cells that were outliers, we performed a PCA using these criteria (Fig. S1d). The PC1 vs PC2 map identified cells that were outliers in with respect to the number and percentage of mapped reads while PC3 identified cells that displayed low number of expressed genes and high levels of mitochondrial reads (Fig. S1d). Cells deemed as outliers were removed from further analysis. After quality filtering, we retained a total of 323 high quality cells (163 cells for ALS and 160 cells for the control). Next, we removed lowly expressed genes (counts < 2) that were expressed in less than 10 cells. In summary, we obtained 323 cells that expressed 14774 genes in total with ALS cells expressing on average 7170 genes while the control dataset expressed 7745 genes with the overall distribution of the number of genes being similar between the two datasets (Fig. S1e). Each gene was classified based on whether it was protein coding, long non-coding, pseudogene or small nuclear/nucleolar RNA. We did not find any systematic difference in distribution of the gene classes expressed between the ALS and controls datasets (Fig. S1f). Finally, principal component analysis using all expressed genes confirmed that our data did not show any batch effects arising from the independent differentiations or the capture plates (Fig. S1g). In summary, our single cell transcriptomes for the ALS and controls sets were similar in quality and character on a genome-wide level. Our differentiation protocol was designed to generate spinal MN[27]. *In vivo*, spinal MN at different rostro-caudal levels of the spinal cord are demarcated by specific combinations of HOX gene expression (known as the HOX code)[28]. To ascertain the rostrocaudal address of our *in vitro* differentiated neurons, we estimated the percentage of cells expressing each of the 39 HOX genes and plotted the data as a heatmap (Fig. 1d). The heatmap showed most of the cells expressed HOXA5 and HOXB5 with very few cells expressing HOXB8 and HOXD8 and none expressing HOX genes from paralog groups 9 and higher (Fig. 1d). This clearly indicated that all of our cells were largely restricted to the hindbrain or brachial spinal cord identity as would be expected from our differentiation protocol that employed 0.1uM retinoic acid without any GDF11[29]. Next, we assessed expression of the classical markers of neuronal subtypes for motor neurons (MNX1, CHAT, PRPH, SLC5A7, ALDH1A2, ACHE, VAChT or SLC18A3), interneurons (GAD1, GAD2, ZFHX2, ZFHX4, LMO3, LMO4, ARX) and non-neuronal cells (PAX6, REST, MKI67, S100B, SOX9, SLC1A3) across all cells (Fig.1e). Our data indicates that iPSC-derived neuronal cultures display wide variation in expression across individual single cells that typically gets averaged in bulk analysis[30]. To resolve this heterogeneity and enable differential expression between relevant classes of neurons, we sought to classify cells into specific neural lineages.

### Classification of single cells into neural subtypes

Gene expression of selective markers in single cells is routinely used to classify neurons into distinct lineages[31, 32]. However, using one or two markers to classify single cells can lead to erroneous classification as single cell data typically has a high rate of dropouts, especially for lowly expressed transcripts. On the other hand, clustering of single cell data based on the expression of all detected transcripts might lead to sub-optimal clustering due to the inclusion of genes that are irrelevant to the classification. We sought to circumvent these problems by first identifying genes that can be used to classify cells into relevant cell types (neurons vs glia and MN vs IN). These lists of genes were termed as classifier gene sets. Cells were classified into distinct neural subtypes by clustering single cells based on the relevant classifier gene set as described below.

#### Neuron vs glia classifier gene set

We first identified genes differentially expressed between neurons and glia using a recently published gene expression dataset on purified human neurons, astrocytes and oligodendrocytes from frozen brain tissue[33]. Genes that displayed a fold change of at least 20 between neurons and astrocytes or oligodendrocytes were included for future analysis. Differential gene expression analysis identified 707 genes as differentially regulated between neurons versus glial cells. Gene ontology using DAVID[34] on the differentially expressed gene set showed enrichment of specific functional categories related to neuronal physiology. Categories related to neuron development such as GABAergic synapse and postsynaptic cell membrane were found to be enriched in the neuron-activated genes while cell cycle and glial differentiation terms were deemed to be enriched in the downregulated genes confirming that our identified gene set was able to distinguish neurons and glia (Fig. S2). Out of the 707 genes, 682 genes were expressed in our filtered single cell dataset. This list of genes was termed neuron_vs_nonneuron.

#### MN vs IN classifier gene set

To identify genesets that can differentiate between MNs and INs, we used the “knowledge matrix” defined previously to classify neurons isolated from embryonic mouse spinal cord into specific subtypes [35]. This a binary matrix mined from published literature on key genes expressed in specific neuronal subtypes. We updated this matrix to include the following genes that are highly expressed in glial cells and progenitors: S100B, SOX9, PAX6, MKI67 and REST (Supplementary data file S2). In total, the matrix comprised of 52 genes and 14 cell types. Next, we extracted the expression data for the 52 genes from our single cell read counts for the healthy cells and binarized the counts for each gene using K-means clustering. We performed this analysis only on the healthy cells to avoid including any influence of the SOD1 mutation on marker gene expression. Our goal was to use this initial set of 52 genes to extract a wider gene set that can be used to classify the ALS and healthy data sets. The expression profile for each gene across the healthy cells was clustered using k=2 that separated the profile into low and high expression clusters. The high expression cluster was used to estimate a threshold for binarizing the gene expression vector (Methods). This resulted in a binary profile for the 52 markers gene for each cell. Each cell was now compared with each of the cell types in the knowledge matrix by estimating a jaccard coefficient (JC), which was used to classify a cell as a MN or IN. Due to the low number of cells, we did not distinguish between IN subtypes at this stage. The top 25 cells (based on the JC) classified as MN or IN were used to generate a differential expression list that could be used to distinguish MNs from INs based on their expression profile. This set of 600 genes, termed the MN_vs_IN geneset, was combined with our neuron_vs_nonneuron list described above. After filtering for genes affected by ambient RNA expression (Methods), we generated a list of 1060 unique classifier genes.

#### Clustering all cells into neural subtypes

We performed unsupervised clustering of all single cells (healthy and SOD1) using our classifier geneset into nine clusters using uniform manifold approximation and projection (UMAP) as part of the Monocle3 package[36, 37] (Fig.1f). Analysis of median expression of known MN, IN and non-neuronal marker genes confirmed that the clustering had successfully grouped cells together by subtypes i.e. MNs(clusters 1,2,3,4), INs (clusters 5,6,7) and non-neuronal glial progenitors (clusters 8,9) (Fig.1g). To evaluate the possibility of mixed clusters i.e. clusters with cells that display expression patterns of multiple cell types, we performed partition analysis that groups clusters together into “super-clusters”. Partition analysis grouped cluster 9 (IN cluster) with the non-neuronal clusters 3 and 4. Similarly the MN cluster 7 was grouped together with the IN clusters 1,2 (Fig.1h). Hence, we termed cells in clusters 9 and 7 as mixed (Fig. 1i). The mixed cells were removed from further analysis. In summary, our clustering approach identified 61 MNs and 41 INs in the healthy dataset, while 38 MNs and 49 INs were identified in the SOD1 ALS dataset (Fig.1j).

### Differential expression analysis of ALS and control neurons

To gain deeper insights into the transcriptional changes within ALS neurons, we compared gene expression in SOD1 ALS MN with the isogenic control MN using DESeq with parameters recommended for single cell RNA-seq (Methods). Differential expression analysis identified 495 upregulated genes and 170 downregulated genes in ALS MN at a p-value threshold < 0.01 (adjusted for multiple hypothesis correction) and fold change > 2 (Fig.2a). On the other hand, comparative analysis between ALS and control IN revealed far fewer genes dysregulated in ALS IN compared to ALS MN at the same threshold (63 genes upregulated while 46 genes downregulated) (Fig.2b, 2c). Additionally, there was minimal, though significant, overlap in the gene expression programs perturbed between the two classes of neurons with 30 genes shared in the upregulated set and 4 genes shared in the downregulated set (Fig. 2d). Further analysis of these genesets revealed that differentially expressed genes in INs had a similar distribution for the fold changes but most genes did not pass the p-value threshold (Fig.S3). This could be due the possibility of multiple interneuron subtypes present in the population resulting in higher gene expression variability. Hence, we first decided to focus on MNs for further analysis.

**Fig. 2.**
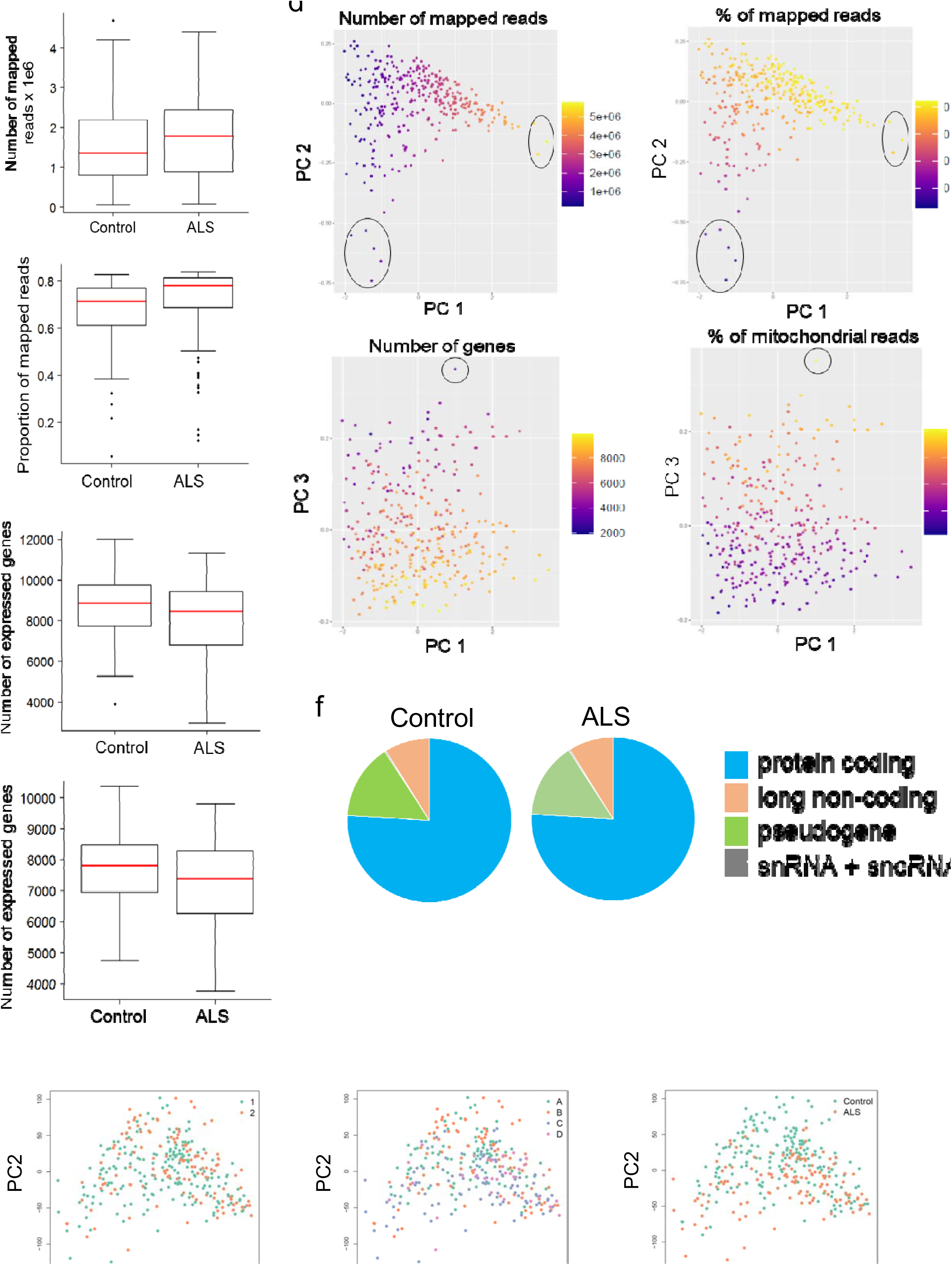
Differential expression analysis of ALS MN and IN. a,b) Heatmap showing expression data for differentially expressed genes between (a) ALS and Control MN, (b) ALS and Control IN. Rows correspond to genes while columns correspond to cells. Normalized log transformed read counts were mean centred for plotting. Clustering of genes and cells was based on the Pearson correlation between rows and columns, respectively. c) Number of genes up or down regulated in ALS MN (blue) compared to ALS IN (orange). d) Overlap between genes up or downregulated between ALS MN and IN. Significance was estimated using the hypergeometric distribution. e) Pathways identified using gene set enrichment analysis (GSEA) on genes up (left panel) or down (right panel) regulated in ALS MN. p-values were adjusted using the Benjamini Hochberg procedure. f) Enrichment analysis of likely pathogenic variants associated with different diseases in genes upregulated in ALS MN. Vertical axis shows terms used to search the ClinVar database to find associated pathogenic variants. ALS: Amyotrophic Lateral Sclerosis, MN: Motor neuron, AD: Alzheimers disease, PD: Parkinsons disease, HD: Huntingtons disease, CMT: Charcot Marie Tooth, SCA: Spinocerebellar Ataxia, MS: Multiple Sclerosis, NDD: Neurodevelopment disorder. The red dashed line indicates a p-value threshold of 0.05.

We performed gene set enrichment analysis (GSEA) to identify pathways enriched in the differentially expressed genesets. This analysis avoids setting strict thresholds and works on the entire geneset that is sorted from most upregulated to most downregulated. GSEA identified several pathways dysregulated in ALS MN. The downregulated gene ontology terms were distributed across 2 main categories, 1) synaptic function and 2) mitochondrial structure and function. Terms enriched related to synaptic function such as “trans-synaptic signalling” and “anterograde trans-synaptic signalling” while mitochondria related terms included “respiratory electron transport” and “oxidative phosphorylation” (Fig. 2e). Additionally, genes involved in calcium homeostasis were also found to be downregulated in SOD1 MNs. On the other hand, terms enriched in the upregulated genes included terms related to the mitotic cell cycle, protein targeting to the endoplasmic reticulum (ER) and RNA splicing (Fig. 2e). Deficiencies in axonal transport, synaptic signalling, mitochondrial oxidative phosphorylation pathway, defects in calcium homeostasis, endoplasmic reticulum stress and increase in cell cycle related genes has been identified previously in bulk analysis of ALS MNs including SOD1 ALS [8, 25, 38–41], which was recapitulated in our single cell expression analysis. Importantly, dysregulation of synaptic signalling and structure in our ALS model supports the dying back hypothesis of ALS that posits neuronal degeneration is secondary to pathology initiated at the distal end of the axon and neuromuscular junction[42]. Finally, we observed that pathogenic variants associated with ALS in the ClinVar[43] database were significantly enriched in genes found to be upregulated in ALS MNs (Fig. 2f). Overall, this analysis showed that our single cell data was able to recapitulate gene expression changes known to be associated with ALS MNs.

### Co-regulated gene modules dysregulated in ALS compared to healthy neurons

To gain a systems level understanding of the observed transcriptional changes, we decided to perform weighted gene co-expression network analysis (WGCNA)[44, 45] to identify modules of co-regulated genes dysregulated in ALS MN and IN. WGCNA identifies sets of genes that are highly correlated (called gene modules) and links these modules with specific phenotypic traits associated with each sample. Since co-expression analysis works on identifying co-regulated sets of genes, a large sample set with high variability per gene across the samples is required for robust network analysis. However, since the single cell data is highly sparse, performing correlation analysis on the read counts might lead to spurious associations or false negatives due to gene dropouts. To overcome this shortcoming, we implemented a recently described k-NN algorithm to smooth the read counts[17] (Methods). Further, we only retained genes that were expressed in atleast 20 cells. The smoothened normalized count data across 189 neurons and 14,054 genes was used to construct modules using topological overlap. WGCNA identified a total of 26 gene modules with sizes ranging from 81 to 1444 genes with a median of 449 genes (Fig. 3a). We used the module eigengene (which is the first principal component for all genes in a given module) as a representative expression of that module in a cell. By estimating the Pearson correlations of each module eigengene with cell type or genotype, we identified modules that were positively or negatively associated with MNs, INs or ALS.

**Fig. 3.**
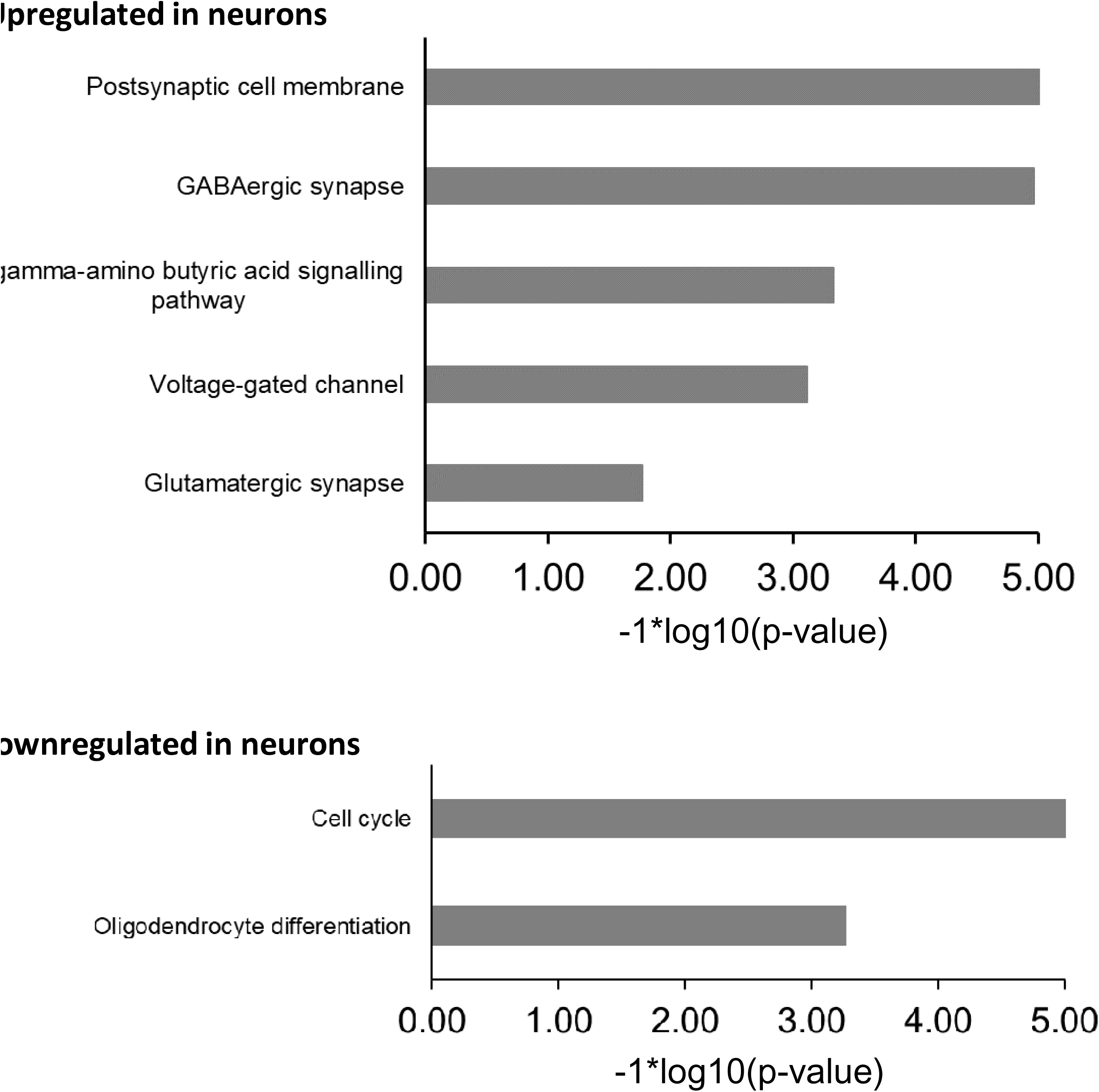
Network analysis of all neurons using Weighted Gene Co-expression Network Analysis (WGCNA) a) WGCNA identified 26 modules of co-regulated genes across the neuronal dataset. Each module was identified by a specific colour. Height indicates the dissimilarity between genes. Dissimilarity was based on the topological overlap between genes. b) Module eigengenes for each module were compared between two groups of cells i.e. ALS MN vs Control (CTR) MN (lower panel) or Control MN vs IN (upper panel). Association of a given module with a specific group was estimated using Pearson correlations. For a comparison of group A vs group B, a correlation of 1 (red) indicates that the module eigengenes were higher in group A compared to group B. A correlation of −1 (blue) indicates that the module eigengenes were lower in group A compared to group B. Correlations with an adjusted p-value < 0.01 were considered significant (outlined in yellow). c) Gene ontology enrichment analysis of modules significantly correlated (positive or negative) with ALS MN. P-values shown were adjusted using the Benjamini-Hochberg procedure. Module lightcyan1 did not yield any significant GO terms and has not been shown. The red dashed line indicates a p-value threshold of 0.05.

Six modules (darkgrey, turquoise, darkslateblue, lightcyan1, darkorange2, pink) were positively correlated while one module (royal blue) was negatively correlated to ALS MN compared to isogenic control MN (adjusted p-value < 0.01 & absolute correlation > 0.4) (Fig. 3b, Fig. S4). GO enrichment analysis of these modules revealed association of each module with specific functional categories. Genes assigned to the royalblue module that was downregulated in ALS MNs were significantly enriched for functional categories related to synaptic function and signalling, axon structure, autophagy and respiratory electron transport (Fig. 3c). These observations were in accordance with the downregulation of these functional categories observed in the differential gene expression analysis of ALS MN. The turquoise module was enriched in genes associated with mitotic cell cycle, TP53 activation and chromatin remodelling (Fig. 3c). Re-entry of post-mitotic neurons into the cell cycle is associated with apoptosis[46], and suggests a possible mechanism of neuronal cell death in ALS. The darkslateblue module was enriched in terms related to RNA processing including translation, splicing and decay. RNA processing defects have been commonly observed in ALS MN[47] while genes involved in translation have been found to display increased expression in sporadic ALS MNs[48]. The pink module was enriched in the terms “coagulation” and “Signalling by Wnt”. Wnt signalling has been found to be activated in bulk analysis of SOD1 ALS MN[25]. The darkgrey module showed enrichment of terms related to catalytic activity. Many of these genes were chromatin remodelling enzymes and these genes may be related to chromatin organization.

Interestingly, the royalblue module associated significantly with healthy MN compared to healthy IN (Fig. 3b), indicating that genes involved in synaptic function and respiration are highly expressed in MNs compared to INs, in accordance with the high metabolic demands on MNs. Downregulation of these genes specifically in ALS MN suggests a possible mechanism that can explain the dysregulation of synaptic functions in ALS MN.

### Pseudotime analysis of degenerating MNs

The dying back model of neurodegeneration posits that disassembly of the neuromuscular synapse and axon retraction are the leading events that precede cell death[42]. Our single cell data provided an opportunity to test whether the observed dysregulated pathways displayed a temporal dependence. Genes differentially expressed in ALS MNs can be considered as a molecular phenotype that drives the cellular phenotype of neuronal degeneration. We used the ALS MN molecular phenotype to pseudotemporally order MNs along a trajectory of degeneration. We first clustered both ALS and healthy MN in a reduced dimensional space using PCA, and used Slingshot[49] to construct a pseudotemporal trajectory (Fig. 4a). Slingshot identified a trajectory that extended from the healthy MNs into the ALS MNs (Fig. 4b). We first confirmed that the cell ordering was not biased by any batch effect (Fig. 4c). Next, we used loess smoothing to model gene expression counts for the differentially expressed genes along the trajectory. Clustering the loess smoothened profiles showed that distinct sets of genes are activated and downregulated in a temporal fashion along the cell trajectory (Fig. 4d). We then asked whether the WGCNA identified modules show differential upregulation or downreglation across this trajectory. For each module, we correlated the module eigengene values per cell with the pseudotime trajectory. Our analysis identified four modules that were positively correlated and two modules that were negatively correlated with the pseudotime (absolute correlation threshold > 0.5 and adjusted p-value < 0.01) (Fig. 4e). In accordance to our WGCNA analysis, the royalblue module showed the strongest inverse correlation with the pseudotime and the darkgrey and turquoise modules showed the strongest positive correlations. This analysis also revealed three other modules (darkgreen, bisque4 and sienna3) that were downregulated along the pseudotime (correlation < −0.4, p-value < 0.01). Pathway analysis revealed that these set of genes were enriched for the functional categories “post-synapse”, “pre-synapse”, “synaptic signalling”, “synaptic vesicle”, “voltage-gated ion channel activity”, “axon”, “dendrite”, “transferase complex”, “exocytosis”, “ubiquitin-dependent protein catabolic process”, “cellular respiration”, “Oxidative phosphorylation” and “PI3K-Akt signalling pathway”. This suggested that individual pathways may display differential activation or inhibition along the pseudotime trajectory. We investigated this hypothesis by estimating mean expression profiles for each of these pathways along the trajectory. To estimate mean expression profiles per pathway, the smoothened profile of each gene assigned to a given pathway was mean centred and the expression at each time point in the trajectory averaged across all genes. The resulting profile was a mean normalized expression profile of the pathway along the trajectory. Correlation of these profiles with the pseudotime trajectory uncovered that genes involved in voltage-gated ion activity and synaptic vesicle formation were downregulated earlier in the trajectory as compared to genes involves in mitochondrial respiration (Fig. 4f). This set of genes was also dysregulated prior to genes involves in cell cycle, chromatin remodelling or TP53 activity (Fig. 4f). This indicates that downregulation of synaptic activity is one of the earliest events in SOD1 ALS MN degeneration.

**Fig. 4.**
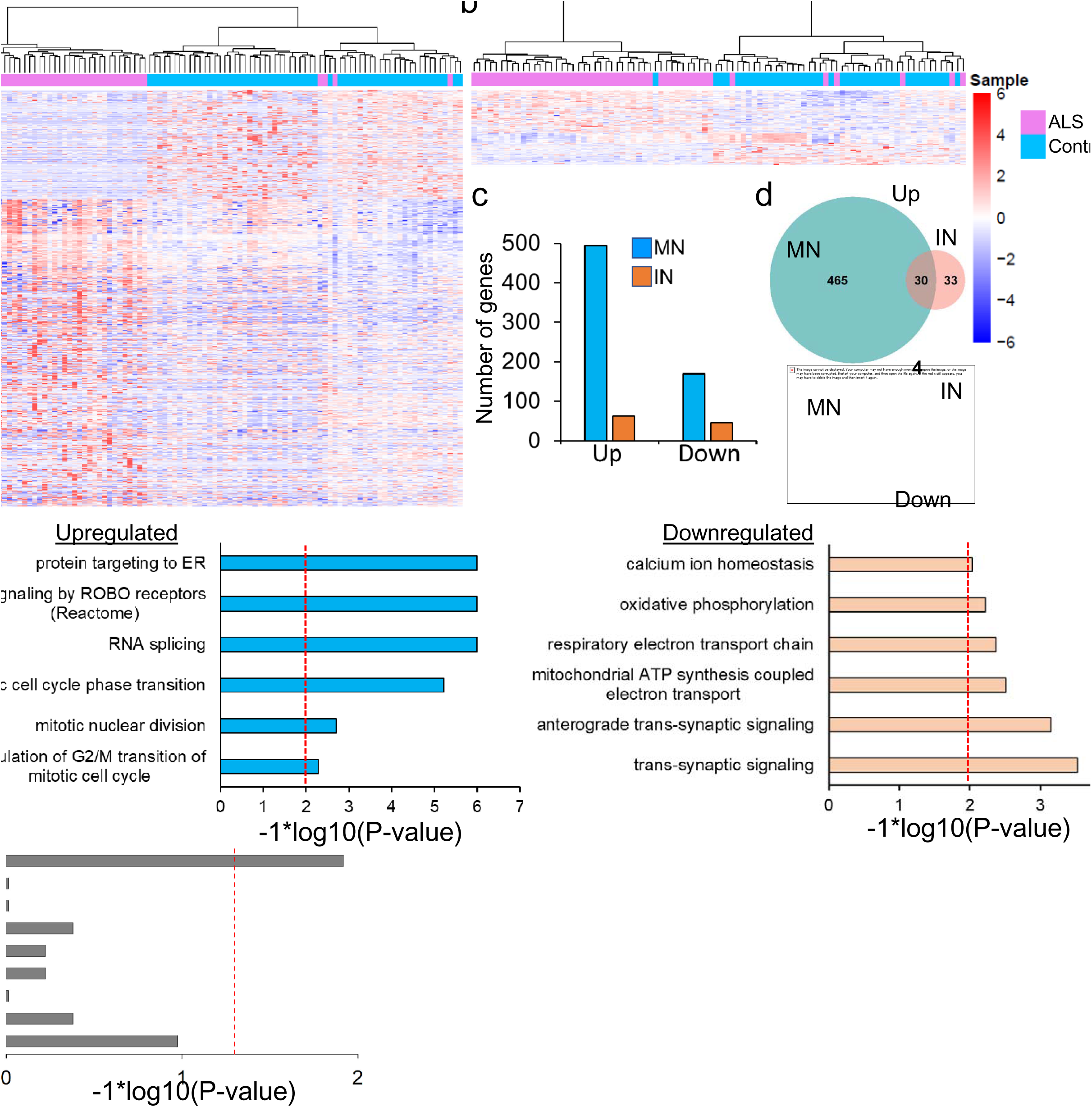
Pseudotime analysis of degenerating MN. a) Pseudotemporal ordering of individual MN. Each dot represents a MN. Each neuron is coloured depending on its position on the pseudotemporal trajectory as shown in the legend on top of the panel. b) Pseudotemporal ordering shows distinct separation of Control and ALS MN. c) Cell trajectory does not show any batch effect. Both differentiation batches were dispersed throughout the plotted trajectory. d) Heatmap of the expression of genes differentially regulated in ALS MN over the pseudotime trajectory. Each gene in this set is plotted along the rows. The log normalized values of each gene were mean centered and smoothened across the trajectory. The smoothened expression values were correlated with the pseudotime and sorted. Genes upregulated over the trajectory are at the top while downregulated genes are at the bottom. e) Heatmap of the expression of WGCNA modules over the pseudotime trajectory. The module eigengene value (first principal component) was used to represent module expression. Eigengene values were mean centered and smoothened across the trajectory. The smoothened eigengene values were correlated with the pseudotime and sorted. f) Heatmap of the expression of signalling pathways enriched in significantly correlated modules across the trajectory. The expression of each gene within a pathway was normalized, mean centered and smoothened as in d. The smoothened expression values across all genes within a pathway were averaged at each time point in the trajectory. The average expression values for each pathway were correlated with the pseudotime and sorted. Pearson correlations were used in panels d,e,f. pt indicates pseudotime.

### Master regulator analysis using single-cell transcriptomics

Having identified a molecular phenotype in ALS MNs, we wanted to identify transcriptional factors (TFs) that were main driver of this phenotype. Identification of such master regulators requires a context-specific transcriptional network[50]. However, building such networks typically requires hundred of samples. Our single cell RNA-Seq data offered a unique opportunity to build such a network as each cell can be considered a distinct biological entity and hence an independent “sample”. Additionally, this network would have the advantage of being context-relevant as the expression data used to build the network would be derived from ALS neurons.

To this end, we deployed the ARACNE algorithm that uses an information theoretic approach to infer transcriptional networks [51]. We used the kNN smoothened and normalized count data across all neurons used for the WGCNA as input to ARACNE that was implemented using the RTN package[52]. After pre-filtering genes that displayed minimal change in expression across the single cells, downstream targets were identified for 1137 TFs amongst 12550 genes expressed in the neuronal cells. The threshold p-value for calling edges can be inferred a-priori from the total number of interactions tested. For example, since we were asking ARACNE to evaluate interactions between 1137 TFs and 12550 target genes, a p-value threshold of 5e-8 would result in < 1 (1137*12550*5e-8) false positive edges to be included in the final network. At a p-value threshold of 5e-8, we identified a total of 1,255,493 edges between 1137 TFs and 12550 target genes with an average of 987 targets predicted per TF.

#### Master regulator analysis to identify TFs driving ALS MN dysfunction

We deployed our network analysis to identify master regulators of ALS MN dysfunction (Fig. S5). As described previously, genes differentially expressed between ALS and isogenic control MNs were used to define a molecular phenotype of the disease. Master regulators were identified based on whether there was a statistically significant overlap between the positive and negative regulon of a TF, and the ALS molecular phenotype. In this case, the goal was to identify TFs that are most likely to drive the differential expression in ALS MNs. For a given TF, if the positive regulon was upregulated and the negative regulon was downregulated in ALS MN, the TF was deemed to be activated. If the inverse was true, the TF was deemed to be inhibited. The master regulator analysis (MRA) identified ∼200 TFs at a FDR < 0.01. To further filter the candidate regulators, we checked for concordance between the expression change of a TF and its regulon, and filtered out non-concordant TFs i.e. where the direction of change of the regulon expression and the TF expression is not the same. For example, if the positive regulon of a TF is found to be activated in ALS MNs, then we would expect that the TF is also upregulated. If the TF was not identified as upregulated, it was filtered out of the analysis. We also filtered out TFs that were not found to be differentially expressed in the ALS MNs compared to control. This identified a core set of 81 TFs (58 activated and 23 inhibited) that satisfied the following criteria: 1) the TFs were differentially expressed in ALS MN compared to control, 2) the regulons of these TFs showed significant association (positive or negative) with the ALS gene expression signature, 3) the TF and its regulon expression was concordant (Fig. 5a). The identified master regulators included TP53, HMGB2, TGIF1 and ZFP36L1 as potential drivers of ALS neurodegeneration i.e. the positive regulons of these TFs as well as the TFs were upregulated in ALS MNs. All of these TFs have been implicated as disease drivers in SOD1 ALS validating our approach[25, 50].

**Fig. 5.**
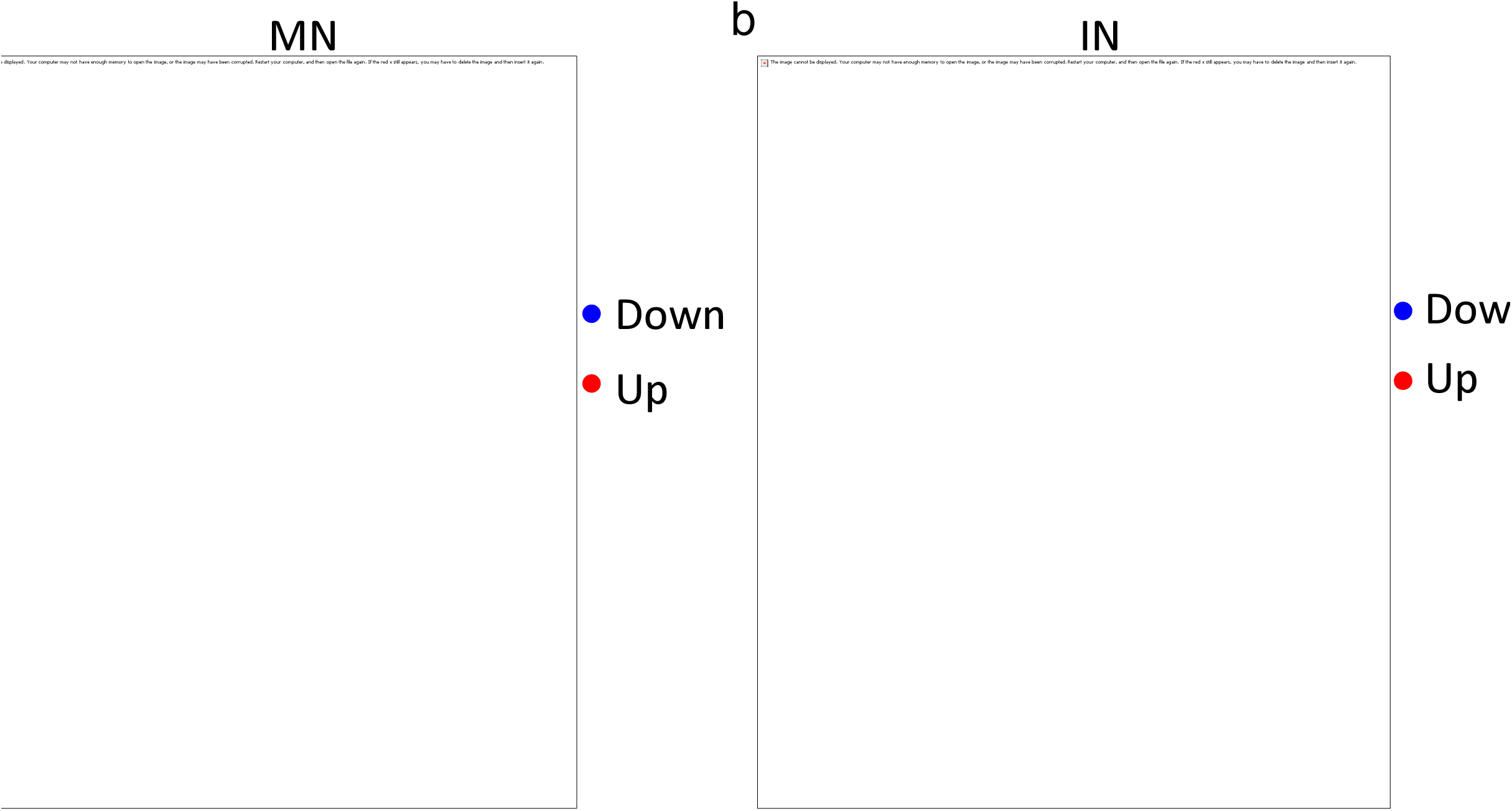
Master regulator analysis (MRA) of ALS disease signature. a) Volcano map showing differential expression of concordant TFs identified by the MRA as the most significant drivers of the ALS molecular phenotype. Each dot is a TF. Red indicates that the TF was significantly upregulated while blue indicates that the TF was downregulated in ALS MN. Black indicates no significant change of expression in ALS MNs. TFs previously found to be associated with ALS MN degeneration are highlighted in grey circles (TGIF1, HMGB2, TP53, ZFP36L1). b) Clustering of master regulators inhibited in ALS MNs based on their mutual information (MI) scores. Higher MI score indicates co-regulation between TFs i.e. the TFs are expressed in a similar manner across the cells, and is highlighted blue. Red square outlines a cluster of TFs that are highly co-regulated. c) Clustering of master regulators activated in ALS MNs based on their MI scores. Higher MI scores are indicated in blue. Red square outlines a cluster of TFs that included TGIF1, ZFP36L1 and HMGB2 as well as mediators of the TGFβ pathway (SMAD2, TGIF2). d) Overlap analysis between regulons and WGCNA modules. Left panel: positive regulons of the TFs highlighted in red in panel (c) were compared with gene modules identified by WGCNA. Significance of the overlap was estimated using a hypergeometric distribution and the p-values were plotted as a heatmap. Right panel: same analysis performed using the negative regulons of the displayed TFs. e) Expression profiles of a subset of TFs identified in panel d that changed across the MN pseudotime trajectory. Each dot represents the expression of the TF within a single MN. Each dot has been coloured based on the position of that cell within the trajectory. Y-axis indicates log2 transformed and normalized expression values. X-axis shows the pseudotime. The black line indicates the loess regressed expression profile of the TF across the trajectory. f) Left panel: Heatmap showing the enrichment of the positive regulons of the master regulator TFs in publicly available ALS datasets. Bulk SOD1 E100G: SOD1 E100G iPSC-derived MN analysed in bulk. GSE54409: SOD1 A4V iPSC-derived MN purified via flow sorting. GSE46298-129Sv: MN laser-capture micro-dissected from SOD1 G93A mouse model, fast progressing strain 129Sv. A: pre-symptomatic, B: onset, C: symptomatic, D: end-stage. GSE46298-C57: MN laser-capture micro-dissected from SOD1 G93A mouse model, slow progressing strain C57. A: pre-symptomatic, B: onset, C: symptomatic, D: end-stage. GSE76220: MN laser-capture micro-dissected from sporadic ALS spinal lumbar tissue. GSE76220-PC1: GSE76220 data was filtered for genes involved in wound healing. GSE18920: MN laser-capture micro-dissected from sporadic ALS spinal lumbar tissue. Enrichment of the positive regulons of the TFs were used as input genesets for a GSEA performed on each dataset. Log transformed P-values were assigned the same sign as the GSEA enrichment scores and plotted as a heatmap. Thus, green indicates that the regulon showed a significant positive enrichment while yellow indicates that the regulon showed a significant negative enrichment in the queried dataset of differentially expressed genes. Right panel: same analysis as the left panel using the negative regulons of the TFs. g) Enrichment of the TGFβ signalling pathway in publicly available ALS MN datasets estimated by performing GSEA. TGFβ pathway datasets were derived from the MSigDB hallmark and KEGG databases, and were used as the input geneset. The red dashed line indicates a p-value threshold of 0.05.

To furter investigate these regulators, we extracted sub-networks of the activated and inhibited TFs, and clustered them on the basis of their mutual information scores. The inhibited TFs formed two clusters where the second cluster (highlighted in red) showed higher co-regulation compared to the first cluster, indicating that these TFs functioned in similar pathways. This cluster included two HOX genes, HOXA1 and HOXD8. HOX genes have important roles in defining MN identity during development[28]. Although, HOX genes have been found to be expressed in adult human and mouse MN[53], their function in MN post-specification is unclear. We considered the possibility that downregulation of HOX genes could be due to specific loss of a MN subtype. For example, loss of hindbrain MN arising from rhombomere 1 would result in downregulation of HOXA1 in the differential expression analysis. To address this possibility, we decided to sub-cluster the MN according to the expression levels of specific HOX genes and then perform differential gene expression analysis for the sub-clusters. Since, HOXA1 was found to be more significantly downregulated than HOXD8 (30 fold vs 8 fold, respectively), we decided to sub-cluster cells based on HOXA1 expression. In the ALS group, 35 out of 38 MN displayed low levels of HOXA1 expression i.e. read count less than or equal to 2. In the isogenic control group 28 MN had HOXA1 levels <=2 while 33 MN had HOXA1 levels > 2. We divided the MN into two groups: the “high” group displayed HOXA1 levels > 2, while the low group displayed HOXA1 levels <= 2. We first compared the ALS and Control MN in the low group. Differential expression analysis revealed 136 genes upregulated and 65 genes downregulated in the ALS MN in the low group (adjusted p-value < 0.01). However, 115 out of the 136 genes were also found in the original 495 genes that were identified without any sub-clustering i.e. comparing all ALS MN with all control MN (Fig. S6a). For the downregulated genes, we found 49 out of the 65 genes were present in the original set of downregulated genes (Fig. S6a). These overlaps were highly significant, indicating that the sub-clusters behaved very similar to the parent set of MN. We further performed GSEA to identify GO terms enriched in the set of differentially expressed genes in the low MN group. GSEA identified exactly the same or very similar terms to be up and down regulated as found when comparing all MN (Fig. S6b). For example, genes related to cell cycle, RNA processing, translation and chromatin were upregulated while genes related to synaptic transmission were downregulated. Next, we wanted to perform a similar analysis for the MN expressing high levels of HOXA1. However, as only 3 ALS MN expressed high levels of HOXA1, we could not perform this analysis. As a proxy, we compared the “low” ALS MN with “high” Control MN. This analysis generated similar results to the previous one (Fig. S6c,d). Finally, we compared Control MN expressing high HOXA1 levels with Control MN expressing low HOXA1 levels to identify subtype-specific effects, if any. Differential expression analysis identified only 2 upregulated genes and 0 downregulated genes at an adjusted p-value threshold of 0.01. As expected, one of the upregulated genes was HOXA1. GSEA identified “memory” as the only GO term upregulated at an adjusted p-value equal to 0.022. Although we cannot completely rule out the possibility, our results indicate that observed downregulation of HOXA1 is not due to loss of a specific MN subtype. This cluster also included the TFs MNX1 and ONECUT1, which are known to be involved in MN homeostasis (Fig. 5b). This suggested that master regulators of MN homeostasis may be downregulated in degenerating ALS MNs. However, we noted that MNX1 was lowly expressed in our dataset (read counts were <=3 in 91 out of the 99 MN), which can exaggerate the fold changes identified by DESeq2. Nevertheless, our data indicates that MNX1 should be used with caution as a marker to identify MN while performing survival analyses.

For the activated TFs, clustering analysis broadly identified four clusters (Fig. 5c). The second cluster (highlighted in red) included the TF SMAD2, a key mediator of the TGF-β signalling pathway. This cluster also included other TFs known to be involved in TGF-β signalling (TGIF1, TGIF2), their downstream targets (ZFP36L1, SOX2) as well as EZH2, a member of the PRC2 complex that works synergistically with the TGF-β pathway[54–57]. TGF-β signalling has been previously observed to be upregulated on spinal astrocytes and muscle of transgenic SOD1 mouse models of ALS[58–60]. Hence, we decided to focus on the set of 18 TFs identified in cluster 2.

We asked whether the identified TFs regulated genes within the WGCNA modules that were significantly associated with ALS MNs. We overlapped the positive and negative regulons of each of the 81 TFs with genes assigned to each module and estimated the significance of the overlap using a hypergeometric test (Fig. S7). We found several TFs whose targets showed significant overlap with the 7 modules (darkgrey, turquoise, darkslateblue, lightcyan1, darkorange2, pink and royalblue) associated with ALS gene expression (Fig. S7). Out of the 18 TFs in cluster 2 in Fig. 5c, the positive regulons of 15 TFs showed a significant overlap with the top 3 modules (darkgrey, turquoise and darkslateblue) deemed to be upregulated in ALS MN (Fig. 5d left panel). On the other hand, the negative regulons of 16 out the 18 TFs significantly overlapped with the royalblue module, which was found to be downregulated in ALS MNs (Fig. 5d right panel). This indicated that the identified master regulators were driving specific gene expression programs in ALS MNs.

Next, we asked whether the identified TFs displayed temporal activation or inhibition across the MN degeneration trajectory. Similar to our earlier pseudotime analysis, we correlated the loess smoothened expression profiles of each TF with the pseudotime and identified 24 TFs as positively correlated and 10 TFs as negatively correlated with the timeline of degeneration (Fig. S8). Overlapping the set of 24 positively correlated TFs with the 18 TFs identified in the second cluster in Fig. 5b highlighted 8 TFs that displayed expression profiles that correlated with MN degeneration (Fig. 5e). We noted that all 8 TFs in this set were significantly associated with the royalblue module while 7 out of these 8 TFs significantly associated with the darkgrey, turquoise or darkslateblue modules (Fig. 5d). Remarkably, 5 out of this 8 TFs (TGIF1, TGIF2, ZFP36L1, SOX2, EZH2) were downstream targets or acted in concert with TGF-β signalling. Though the expression of SMAD2, a key mediator of the TGFβ pathway, was not deemed to correlate with the pseudotime, this is not surprising as the key mode of SMAD2 activation is by nuclear translocation of the phosphorylated protein and not by upregulation of transcript levels. However, correlation of TGFβ activated TFs along the pseudotime provided strong evidence that the TGFβ signalling may be activated in ALS MNs and activation of this pathway correlates with MN degeneration.

Further, we asked whether dysregulation of the identified master regulators is observed in previously published datasets that identified genes differentially expressed in ALS MNs (Fig. S9). We performed GSEA on the published ALS datasets where the positive regulons of the master regulators were used as input genesets for the GSEA. The GSEA assigned a significance score to each regulon based on whether it was deemed to be activated or suppressed in a given ALS dataset. If the positive regulon of a TF was found to be activated in a given dataset, the TF corresponding to the regulon was also deemed to be activated in that dataset. On the other hand, if the positive regulon was found to be inhibited in a given dataset, the TF corresponding to the regulon was also deemed to be inhibited in that dataset. The regulons of most of our activated master regulator TFs (56 out of 58) were found to be activated in at least one ALS dataset (Fig. S9). Regulons of TFs related to MN homeostasis and identity such as MNX1, ONECUT1, HOXA1 and HOXD8 were found to be downregulated in sporadic ALS MN and in the end-stage SOD1 G93A 129Sv MN but not in MN derived from ALS SOD1 iPSC (Fig. S9). This suggested that downregulation of MN homeostatic regulators could be a terminal event in dying neurons.

All 8 TFs identified in the pseudotime analysis (Fig. 5e) were found to be activated in iPSC-derived MN, SOD1 mouse models and sporadic ALS MN (Fig. 5f). Additionally, SMAD2 was found to be activated in all the different models of ALS analysed, including the mouse SOD1 G93A MN at the onset stage (Fig. 5f). These observations led us to hypothesize that the TGFβ signalling pathway is activated in ALS MN. We directly tested this hypothesize by analysing whether genes involved in TGFβ signalling are upregulated in ALS MN. We used the TGFβ signalling datasets from the hallmark collection included in the Molecular Signatures Database (MSigDB), which was designed to reduce noise and redundancy for GSEA[61] as well as the KEGG database[62]. Our analysis confirmed that the TGFβ pathway is indeed activated in SOD1 G93A mouse, SOD1 E100G ALS, SOD1 A4V ALS and sporadic ALS MN (Fig. 5g). Though upregulation of TGFβ signalling has been observed previously in astrocytes and muscle cells derived from the SOD1 G93A ALS mouse model, the activity of TGFβ signalling and the functional consequence of TGFβ activation in ALS MN have not been investigated.

SMAD2 is a downstream effector of the transforming growth factor β (TGFβ) signalling pathway. Upon activation of TGFβ signalling, SMAD2 is phosphorylated and translocates to the nucleus from the cytoplasm. Nuclear SMAD2 regulates downstream gene expression in partnership with other TGFβ mediators such as SMAD4. We assayed levels of phosphorylated SMAD (pSMAD2; the activated version of SMAD2) in ALS SOD1 and isogenic control MNs at day 30. At this stage, ALS MN did not display any significant differences in survival compared to the control MN. Immunostaining assays confirmed that ALS SOD1 MNs display higher levels of nuclear pSMAD2 (Fig.6a). Higher levels of pSMAD2 immunoreactivity have also been observed within motor neuron nuclei in post-mortem spinal tissue derived from sporadic ALS patients[63]. Further, we confirmed upregulation of TGFβ downstream target genes TGFB1, TGFBI and ZFP36L1 (Fig.6b). Upregulation of TGFβ downstream target genes was also observed in MNs differentiated from FUS P525L iPSCs compared with their respective isogenic controls[64] (Fig.S10, Fig.6b). Next, to ascertain whether TGFβ activation contributes to neurodegeneration or is simply a downstream effect of degenerating neurons, we treated SOD1 ALS MNs with the TGFβ inhibitor SB431542. SOD1 MNs were treated at different concentrations of SB431542 starting at day 30 till day 40. Neuronal survival was followed over the course of treatment by counting nuclei at day 30 and day 40. Treatment with SB431542 significantly enhanced neuronal survival in a dose-dependent manner (Fig.6c). Inhibition of TGFβ signalling also decreased apoptotic levels in the SOD1 MNs by day 40 (Fig.6d). Conversely, treatment of SOD1 MNs with TGFβ from day 30 till day 40 resulted in enhanced apoptosis in the neuronal cultures (Fig.6e). In summary, our results demonstrate that activation of TGFβ signalling is a driver of ALS associated MN degeneration and is a shared event between SOD1, FUS and sporadic ALS MNs.

**Fig. 6.**
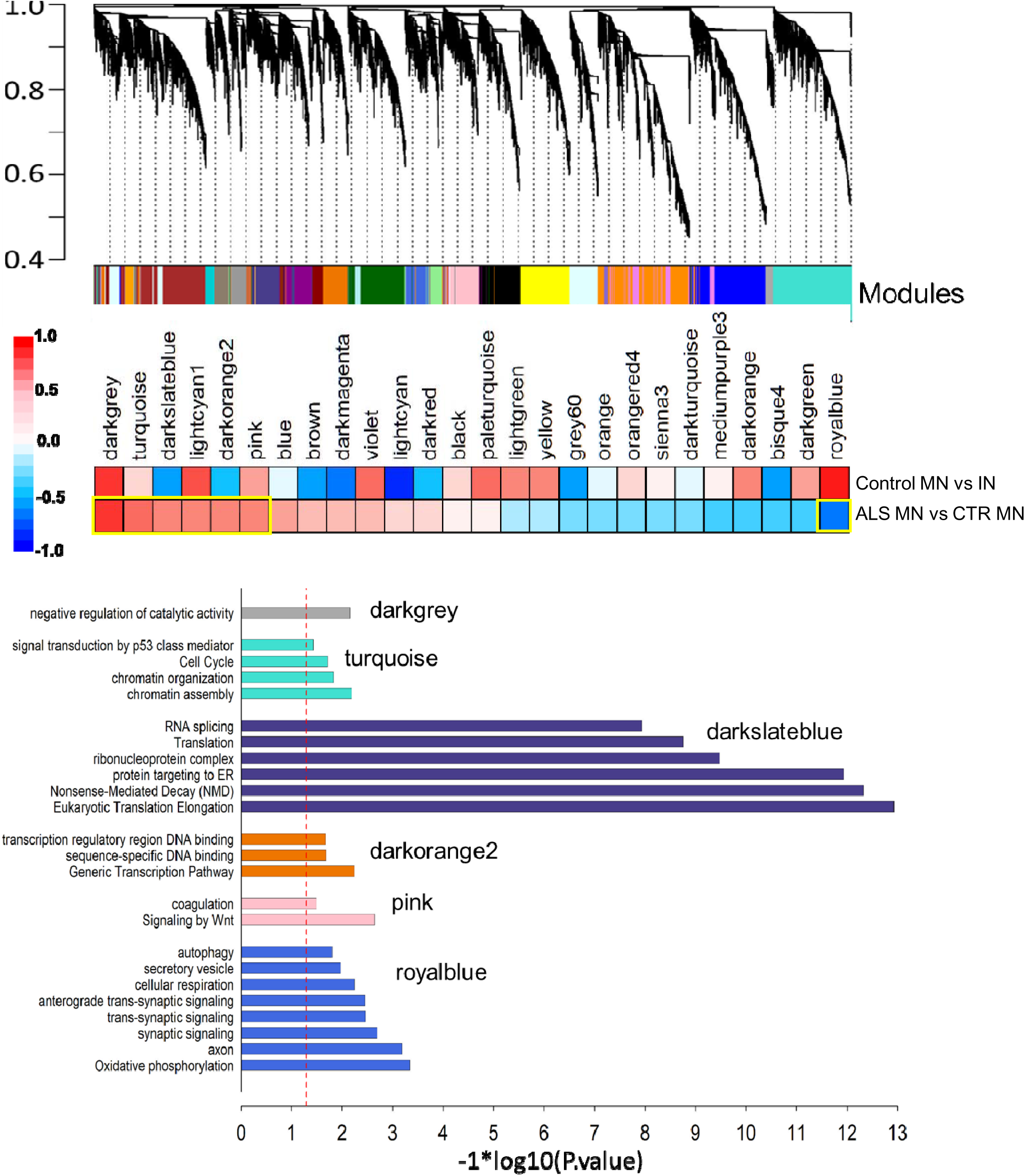
TGFβ signalling is a key driver of ALS MN degeneration. a) Immunofluorescence analysis of phosphorylated SMAD2 (p-SMAD2) in SOD1 ALS (E100G) and isogenic control (E100E) neuronal cultures. Nuclei were stained with Hoechst 33342. Nuclear p-SMAD2 intensities were quantified across three independent differentiations for the ALS and Control cultures. Pyknotic nuclei (white arrows) were excluded from the analysis by setting a size threshold. Scalebar represents 52µm. Median intensity values per nuclei were estimated across at least 100 nuclei per replicate and averaged. Values obtained for each replicate were normalized by the average values in the control neurons. Normalized values were plotted as a bar graph. E100G: ALS SOD1 E100G, E100E: Isogenic control. b) Quantitative RT-PCR analysis of genes upregulated by TGFβ signalling (TGFB1, TGFBI and ZFP36L1). Left panel: Fold changes observed in ALS SOD1 E100G MN, right panel: Fold changes observed in FUS P525L MN. In each case, fold changes were normalized to the respective isogenic controls. c) Quantitation of nuclei in ALS SOD1 MN cultures after treatment with SB431542 at the indicated concentrations. Number of neurons at day 40 were compared relative to day 30. d) Relative annexin V levels in day 40 ALS SOD1 MN after treatment with SB431542 10uM. Annexin V levels were normalized to those obtained in the DMSO only control. e) Relative annexin V levels in day 40 ALS SOD1 MN after treatment with TGFβ 100ng/ml. Annexin V levels were normalized to those obtained in the PBS only control. (n=3 independent differentiations, error bars indicate SEM, **: p < 0.05, *: p < 0.05, p-values were estimated using two-tailed students t-test).

### ALS MNs display re-activation of progenitor expression programs

Network analysis using WGCNA has previously revealed disruption of age-related modules and pathways in sporadic ALS MN[48]. For example, the expression of genes involved in translation was found to decrease with age but was strongly upregulated in sporadic ALS MN. We observed that the module darkslateblue that significantly correlated with ALS SOD1 MN was enriched in genes with a role in translation and RNA processing. This led us to question whether ALS MN displayed aberrant activation of signalling pathways that are active during development. To test this hypothesis, we obtained gene expression profiling data over the time course of motor neuron differentiation from neuromesodermal progenitors (D0) to motor neurons (D15) (GSE140747)[22]. We used DESeq2 to generate a list of genes differentially expressed between D0 neuromesodermal progenitors (NMP) and D15 neurons. This list was sorted such that genes most upregulated in the NMP were at the top while genes most downregulated in the NMP were at the bottom. This process was repeated for time points D1 to D8 generating a set of differentially expressed ranked genelists for neural progenitors. By reversing the genelist D7 vs D15, we generated an additional ranked list of genes that had neuron-specific genes at the top of the list. Additionally, we also generated ranked gene expression profiles in immature (D21) and mature (D35) motor neurons differentiated from healthy iPSCs[23, 24]. Thus, D0 to D8 genelists display progenitor-enriched genes at the top while D15, D21 and D35 genelists display neuron-enriched genes at the top. We used GSEA to quantify the enrichment of each WGCNA module in each genelist (Fig. 7a). In accordance with our expectation, module darkslateblue was significantly enriched in the progenitor genelists D0 to D8, indicating that genes assigned to this module are highly active earlier in motor neuron development. Additionally, modules turquoise and lightcyan1 that were found to be activated in ALS MN were also enriched at earlier developmental time points. Remarkably, genes in module royalblue that was downregulated in ALS MN were enriched in the D15, D21 and D35 neuron-enriched genelists. This indicated that ALS SOD1 MN display activation of genesets that are found to be highly enriched in progenitors compared to mature neurons. These observations were also re-capitulated by analysing the set of differentially expressed genes in ALS SOD1 MN. Genes found to be upregulated in SOD1 MN were significantly enriched in genelists D0 to D8, while downregulated genes were enriched in the neuronal genelists D15, D21, D35 (Fig. 7b). To assay whether these observations hold true for other ALS datasets, we looked for module enrichment in genes upregulated in SOD1 iPSC-derived MN, SOD1 G93A mouse MN and sporadic ALS MN (Fig. 7c). The turquoise module was found to be enriched in the iPSC-derived SOD1 A4V MN, SOD1 mouse model (129Sv) and one of the sporadic ALS datasets (GSE18920). The darkslateblue module was also found to be enriched in the iPSC-derived SOD1 A4V MN and at the onset stage in both SOD1 mouse models (129Sv and C57). Remarkably, the turquoise and darkslateblue modules were found to upregulated in at least one of the sporadic ALS datasets. This enrichment was observed even after removal of genes involved in wound healing for the GSES76220 dataset (Fig. 7c). The royalblue module was found to be significantly downregulated in all the three different models i.e. the SOD1 ALS iPSC-derived MN, mouse SOD1 MN and sporadic ALS MN. Surprisingly, the bulk SOD1 E100G dataset did not show enrichment of the turquoise or darkslateblue modules. This dataset represents gene expression counts derived from bulk RNA-seq analysis without purifying MN from the iPSC-derived neuronal cultures, which could have reduced the sensitivity of detection. Finally, we observed that the positive regulons of several master regulators including TFs activated by the TGFβ pathway (TGIF1, TGIF2, ZFP36L1, SOX2) were highly enriched in the progenitors genelists (Fig. 7d). This indicated that transcriptional mediators of the TGFβ pathway may be responsible, at least in part, for reactivation of these developmental programs. Overall, our results indicate that gene expression programs that are highly active in progenitors are upregulated in ALS MN.

**Fig. 7.**
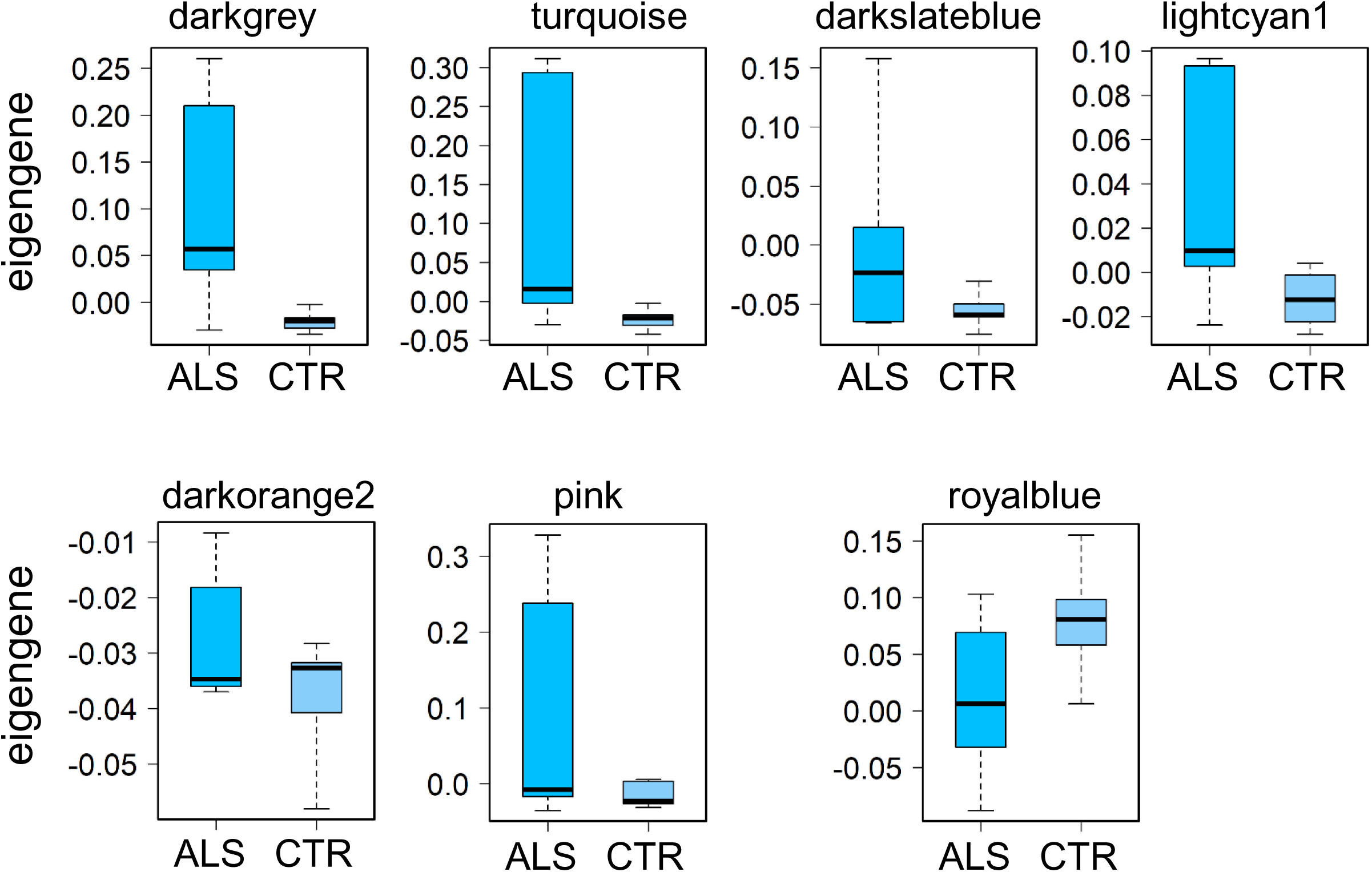
Progenitor gene expression programs are reactivated in ALS MN. a) Enrichment of WGCNA modules in genes upregulated in neural progenitors compared to neurons. Day 0 (D0) indicates neuromesodermal progenitors (NMP). D1 to D8 indicate intermediate time points as the NMP differentiated into MN. Each progenitor stage was compared with D15 MN using DESeq2. Thus, D0 in the panel represents differentially expressed genes between D0 NMP and D15 MN. Accordingly, D1 to D8 represents differentially expressed genes between intermediate progenitors and D15 MN. Differentially expressed genes were sorted such that gene upregulated in progenitors were at the top of the list. To obtain genes upregulated in D15 MN, the gene list obtained from the D7 vs D15 comparison was inverted such that genes upregulated in D15 MN were at the top of the list. D21 and D35 MN genelists were obtained by comparing D21 and D35 MN with iPSC. The red rectangle highlights modules activated in ALS MN while the green rectangle highlights the single module downregulated in ALS MN. Enrichment was estimated using a one-way GSEA where the modules genes were used as genesets and only positive enrichments were calculated. Heatmap shows log transformed p-values. b) Genes upregulated in ALS were significantly enriched in progenitors. GSEA performed was similar to (a) where the genesets used for the GSEA were genes upregulated (Up) or downregulated (Down) in ALS. c) Enrichment of WGCNA modules in publicly available ALS datasets. The datasets are as described in figure 5. Bulk SOD1 E100G: SOD1 E100G iPSC-derived MN analysed in bulk. GSE54409: SOD1 A4V iPSC-derived MN purified via flow sorting. GSE46298-129Sv: MN laser-capture micro-dissected from SOD1 G93A mouse model, fast progressing strain 129Sv. A: pre-symptomatic, B: onset, C: symptomatic, D: end-stage. GSE46298-C57: MN laser-capture micro-dissected from SOD1 G93A mouse model, slow progressing strain C57. A: pre-symptomatic, B: onset, C: symptomatic, D: end-stage. GSE76220: MN laser-capture micro-dissected from sporadic ALS spinal lumbar tissue. GSE76220-PC1: GSE76220 data was filtered for genes involved in wound healing. GSE18920: MN laser-capture micro-dissected from sporadic ALS spinal lumbar tissue. Genes assigned to each module were used as input genesets to the standard GSEA performed on the ALS datasets. The x-axis shows the modules. The red rectangle highlights modules activated in ALS MN while the green rectangle highlights the single module downregulated in ALS MN. Log transformed P-values were assigned the same sign as the GSEA enrichment scores and plotted as a heatmap. Red indicates that the module showed a significant positive enrichment while green indicates that the module showed a significant negative enrichment in the queried dataset. d) Enrichment of the positive regulons of the 81 master regulators identified by the MRA. Enrichment was estimated using a one-way GSEA where the positive regulons of each TF were used as genesets and only positive enrichments were calculated. The differentially expressed sorted progenitor and MN genelists were the same as those used in panel a. Heatmap shows log transformed p-values. Red arrows highlight the 8 TFs identified in Fig 5e that include mediators of the TGFβ signalling pathway.

### ALS V1 interneurons show gene expression changes similar to MN

A recent single-cell transcriptomic study analysing mouse G93A ALS brainstem tissue identified dysregulation of gene expression in spinal interneurons[65]. Hence, we investigated whether mutant SOD1 induces gene expression changes in spinal IN similar to MN. We used the knowledge matrix to classify the 90 IN identified in our single-cell data into 49 V1 (26 SOD1 and 23 healthy), 9 V2a (1 SOD1 and 8 healthy), and 21 V2b (16 SOD1 and 5 healthy) IN. The remaining 11 neurons could not be unequivocally classified into a specific subtype. The V2a and V2b IN numbers were skewed towards the healthy and SOD1 genotypes, respectively. This could probably be due to under-sampling of the IN population while collecting single cells. Hence, we focused on the V1 population for further analysis. Spinal V1 IN are a diverse group of neurons that show highly variable gene expression patterns[66]. The V1 neurons identified in our dataset displayed expression of TFs known to be expressed or enriched in the V1 population[66] (Fig. S11a). Differential expression analysis identified only 2 genes as upregulated and 7 genes to be downregulated in the ALS V1 population (absolute fold change > 2 & adjusted p-value < 0.01) (Fig. S11b). A threshold-independent analysis using GSEA did not reveal any enrichment of pathogenic variants associated with ALS listed in the ClinVar database (Fig. S11c). Surprisingly, genes associated with neurodevelopmental disorders were found to be enriched in ALS V1 IN (Fig. S11c). We performed pathway enrichment analysis using GSEA to avoid using hard thresholds on the differential gene expression data. Interestingly, GSEA identified gene ontology terms related to ER stress, mitotic cell cycle, RNA splicing and translation to be upregulated in ALS V1 IN, similar to those found in ALS MN but with lower enrichment scores (Fig. S11d). Upregulation of translation in ALS V1 IN was also observed in a recent single cell study of sporadic and C9ORF72 iPSC derived neurons[67]. On the other hand, genes involved in synaptic signalling were found to be downregulated, though to a lower extent than observed in ALS MN (Fig. S11d). However, terms related to oxidative phosphorylation or respiratory electron transport that were identified in ALS MN were not observed in genes downregulated in ALS V1 IN, even when the p-value threshold was reduced to 0.1. These observations were recapitulated when we analysed the correlation between WGCNA modules and V1 IN disease status. Out of the 6 modules found to correlate positively with ALS MN, 5 (darkgrey, turquoise, darkslateblue, lightcyan1 and pink) were also identified to correlate positively with ALS V1 IN (Fig.S11e). However, the royalblue module (enriched in genes involved in synaptic signalling and respiration) that was significantly downregulated in ALS MN was not deemed to be perturbed in ALS V1 IN (Fig. S11e). This indicates that genes involved in oxidative phosphorylation are not affected in ALS V1 IN compared to MN.

## Discussion

ALS patient-derived iPSC have provided unprecedented access to human diseased motor neurons enabling researchers to follow the course of degeneration in dish[6]. Investigation of these models using genomics has uncovered key pathways dysregulated in ALS neurons. However, so far, iPSC derived neuronal cultures have been typically analysed in bulk. Hence, it has been difficult to assign observed pathway aberrations specifically to MN or other neuronal subsets present in the *in vitro* culture. The advent of single cell genomics has now allowed the analysis of individual neurons in mixed cultures[30, 68]. This game-changing technology now allows identification of genome-wide gene expression in individual neurons thereby enabling neuronal classification into subtypes. We have applied this technology to analyse RNA expression in individual neurons derived from ALS patient-derived iPSC and the corresponding isogenic controls. This not only allowed us to distinguish gene expression changes in MN and IN, but also enabled us to construct context-specific gene regulatory networks. Though the total number of cells captured in our study is less than typically seen in droplet-based assays, we were able to sequence each cell to a much greater depth (1.5e6 read per cells). This is in contrast to droplet-based studies that result in just 100,000 reads per cell[67]. As a result, our data identified almost thrice the number of genes typically seen in droplet-based experiments. This allowed sensitive identification of differentially expressed genes and pathways. Importantly, it enabled us to map whole transcriptome gene networks. Network analysis is a powerful way to understand how genes interact with each other to bring about cellular phenotypes[69, 70]. By building a transcriptional network specific to the ALS landscape, we were able to associate gene modules with neuronal subtypes specific to ALS or healthy samples. Further, by using master regulator analysis, we were able to identify TFs that drive disease-associated expression changes. Specifically, our analysis identified SMAD2-mediated TGFβ signalling as a key driver of ALS MN degeneration.

The dying back hypothesis posits that neuropathology is initiated in the distal axons and synapses of motor neurons subsequently leading to axon retraction and degeneration of the soma proximally[42]. In support of this hypothesis, defects in the neuromuscular junction and distal axons were identified in mouse models of ALS before onset of symptoms[71]. We have observed similar deficits in axon structure and branching in MN derived from FUS P525L ALS iPSC and in FusΔ14 mouse MN that show severe mislocalization of Fus to the cytoplasm[72]. Additionally, defects in synaptic activity have been observed in MN derived from ALS associated C9ORF72 repeat expansion and mutant TDP43 patient derived iPSC[9, 10]. Interestingly, these defects were observed on prolonged *in vitro* culture of the MN correlating with onset of degenerative phenotypes[9]. Additionally, our analysis indicated that genes involved in synaptic transmission were downregulated earliest in the pseudotemporal trajectory of MN dysfunction. Collectively, these observations strongly indicate that axonal and synaptic dysfunction is an early event that precedes neurodegeneration in ALS. Synaptic collapse can be a downstream effect of either impaired delivery or production of synaptic proteins and mRNAs. Impaired delivery can occur secondary to inefficient axonal transport[73–75]. On the other hand, our data reveals inhibition of the synaptic genes at the transcriptional level thereby impairing production. We find that the observed downregulation of the synaptic program could be due to the dysregulation of master regulator TFs in ALS MNs.

Master regulator analysis identified mediators of the TGFβ signalling pathway to be activated in ALS MNs. Our data confirmed a direct role of TGFβ activation in causing neuronal death in SOD1 ALS MNs. TGFβ activation was also observed in FUS P525L ALS MNs as well as MNs microdissected from sporadic ALS patients. This indicates that an activated TGFβ pathway may be a shared mechanism of neurodegeneration in familial and sporadic ALS MNs. In support of this hypothesis, sporadic ALS MN display elevated levels of phosphorylated SMAD2, the active form of SMAD2, in their nuclei compared to their healthy counterparts[63]. Previous studies have postulated that elevated levels of TGFβ signalling arising from ALS astrocytes or muscle could drive MN death[58–60]. Our study indicates that MN themselves could be the source of TGFβ. Whether astrocytes secrete TGFβ independently or in response to MN remains to be seen.

How does an activated TGFβ contribute to MN dysfunction and death? Analysis of TFs associated with TGFβ signalling provide insights into the underlying mechanism. The negative regulons of SMAD2, TGIF1 and TGIF2 i.e. genes inhibited by these TFs, were enriched for genes involved in synaptic transmission (Fig. S12). On the other hand, the positive regulons of the TFs TGIF2 and ZFP36L1 were enriched for genes involved in mitosis (Fig. S12). These TFs may contribute to MN death by reactivating the cell cycle in post-mitotic neurons. Thus, our analysis identifies a core set of master regulators that drive MN dysfunction and death, by targeting genes involved in synaptic function and cell cycle.

A previous study employing network analysis using WGCNA uncovered disruption of aging networks in sporadic and familial SOD1 ALS MNs[48]. Specifically, gene modules that correlated positively with aged neurons were downregulted in ALS MNs. For example, genes involved in translation and RNA processing decreased in expression over the course of the normal aging process but were observed to be upregulated in sporadic ALS MNs. We observed similar trends in our single cell network analysis where modules showing enrichment of genes involved in translation, RNA decay and splicing were upregulated in ALS MN. This inverse correlation between aging and ALS MN gene exression led us to hypothesize that ALS MN may display reactivation of developmental pathways. Our analysis indicated that gene sets that are highly active in neural progenitors that appear early in MN development are reactivated in ALS MN. Additionally, several of the newly identified master regulators associated with ALS MN gene expression were found to regulate targets that are active in these neural progenitors. Overall, our analysis provides strong evidence to the possibility that developmental gene expression programs may be reactivated in ALS MN. This could also contribute to the observed reactivation of cell cycle and downregulation of synaptic genes in diseased MN, eventually leading to the dismantling of the NMJ. However, whether reactivation of progenitor expression programs are the cause or a downstream effect of TGFβ signalling needs further investigation.

Differential neuronal susceptibility has been recognized in ALS, with oculomotor neurons (OMN) being relatively resistant to neurodegeneration. Laser-capture microdissection followed by gene expression analyses has uncovered distinct gene expression programs active between susceptible spinal MN and resistant OMN[76]. However, whether spinal interneurons are equally susceptible or resistant to neurodegeneration in ALS is unclear. Our results indicate that ALS V1 IN share many of the dysregulated pathways observed in ALS MN. However, genes involved in oxidative phosphorylation or the respiratory electron transport seem to be unaffected in ALS V1 IN while these were found to be significantly downregulated in ALS MN. Given the high metabolic demands of ALS MN, perturbation of mitochondrial pathways could make these neuron more susceptible to degeneration than V1 IN. On the other hand, ALS V1 IN also display upregulation of the same gene expression programs as observed in MN, although to a weaker extent. This would suggest that V1 IN might also display survival deficits but may be relatively more resistant to degeneration than MN.

### Conclusions

The underlying cause of MN degeneration in ALS is very likely to be multi-factorial with multiple drivers collaborating to cause MN demise. We have identified that dysregulation of TFs that disrupt MN homeostasis are major contributors to MN death in ALS. Our results display the power of combining network analysis with single cell trancriptomics to iPSC based neurodegenerative models to uncover drivers of motor neuron degeneration in ALS. We expect that wider use of single cell genomics especially multi-omics technologies to measure different molecular entities from the same cell [77] combined with network biology will help uncover novel regulators that can be targeted using small molecules or gene therapy.

## Availability and Accession Numbers

The raw data has been submitted to ArrayExpress with accession number E-MTAB-7353.

## Acknowledgements

We thank the sequencing centre at the Genome Institute of Singapore for help with Illumina sequencing and mapping. We also thank Peter Henley for his help in formatting the GSE18920 dataset.

## Authors’ contributions

A.B and L.W.S provided funding for the project. A.B. S.C.N, S.H. L.O.G designed and conducted the experiments. A.B., P.T, R.A, C.R.G.W analysed the data. All authors contributed towards interpreting the data and writing the manuscript.

---

**Fig. S1.**
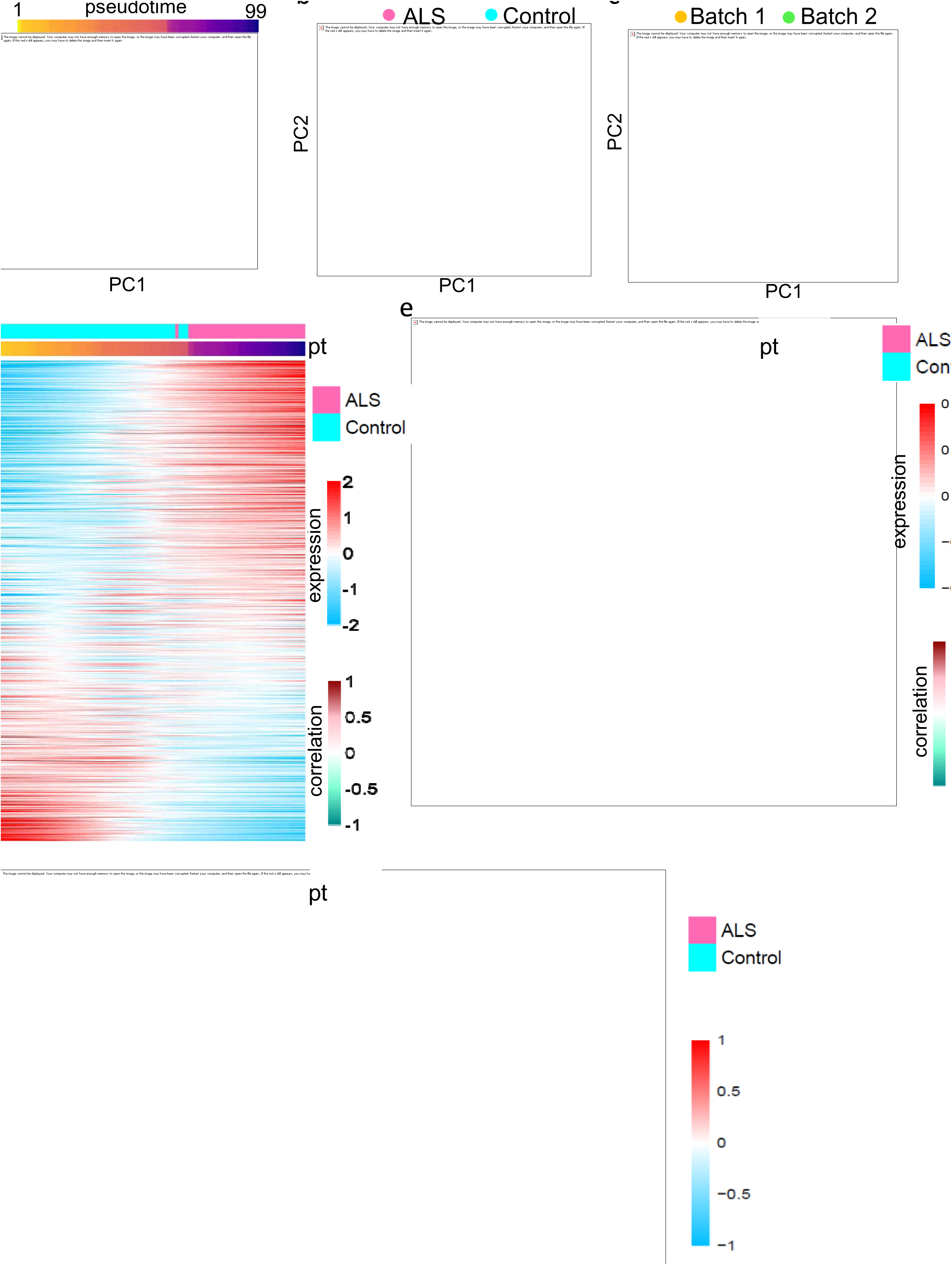
Quality check of the single cell data. a,b,c) Boxplots showing (a) number of mapped reads, (b) proportion of mapped reads, (c) number of expressed genes, in control and ALS datasets before filtering. d) PCA plots displaying the different quality metrics used to filter cells. Each dot represents a cell. The black ellipses mark cells classified as outliers. Overall, nine cells were identified as outliers. e) Boxplots showing number of expressed genes, in control and ALS datasets after removing outlier cells. f) Pie chart showing the distribution of the expressed genes in different classes. g) PCA plots generated using all expressed genes show there is no batch effect between the datasets. Each dot represents a cell. Left panel: cells have been coloured based on the replicates. Middle panel: cells have been coloured based on which C1 fluidigm plate they were captured in. A: Control replicate 1, B: Control replicate 2, C: ALS replicate 1, D: ALS replicate 2. Right panel: Cells have been coloured based on genotype (i.e, isogenic control or ALS SOD1 E100G).

**Fig. S2.**
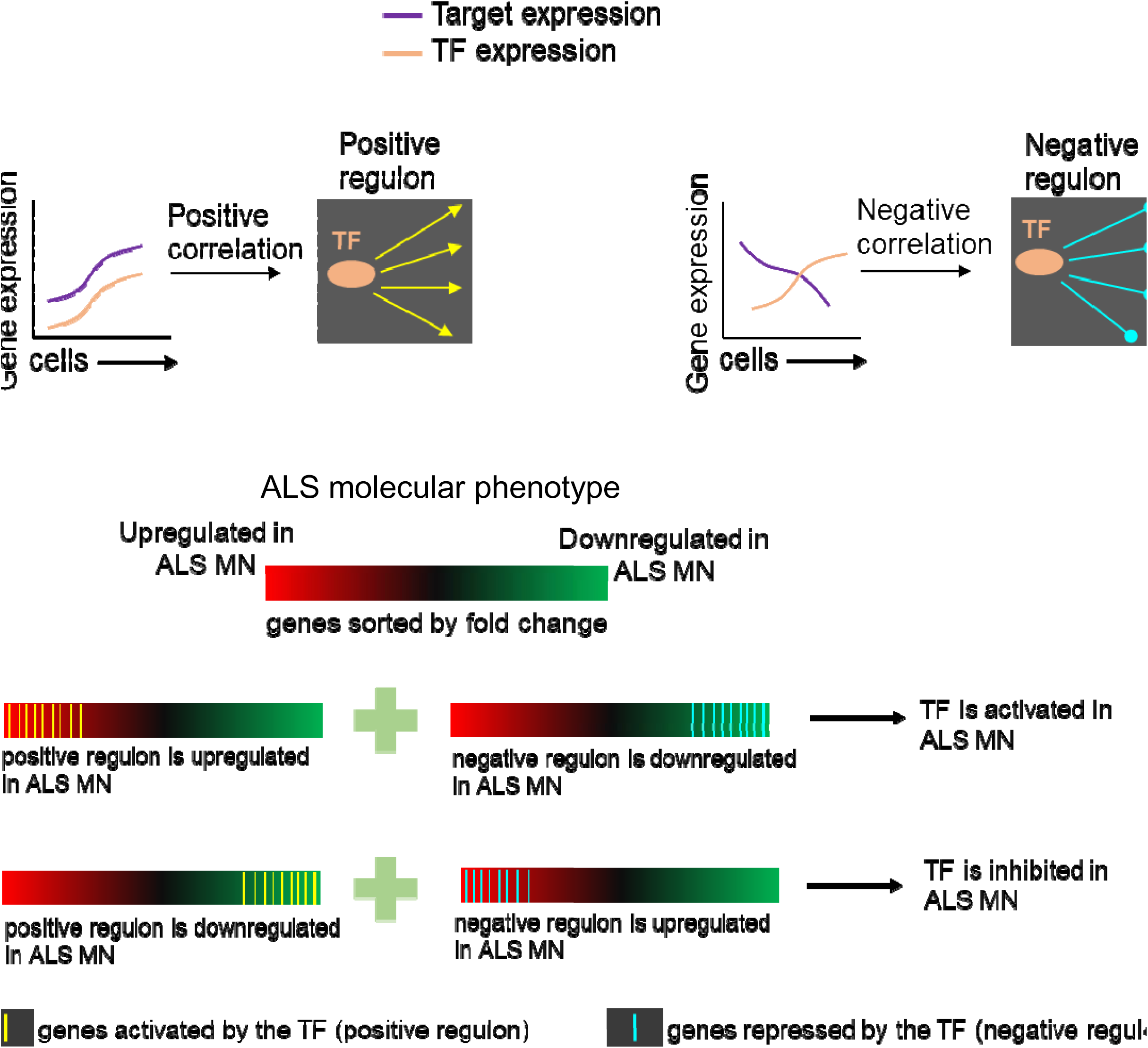
GO enrichment analysis of neuronal vs glial classifier gene set.

**Fig. S3.**
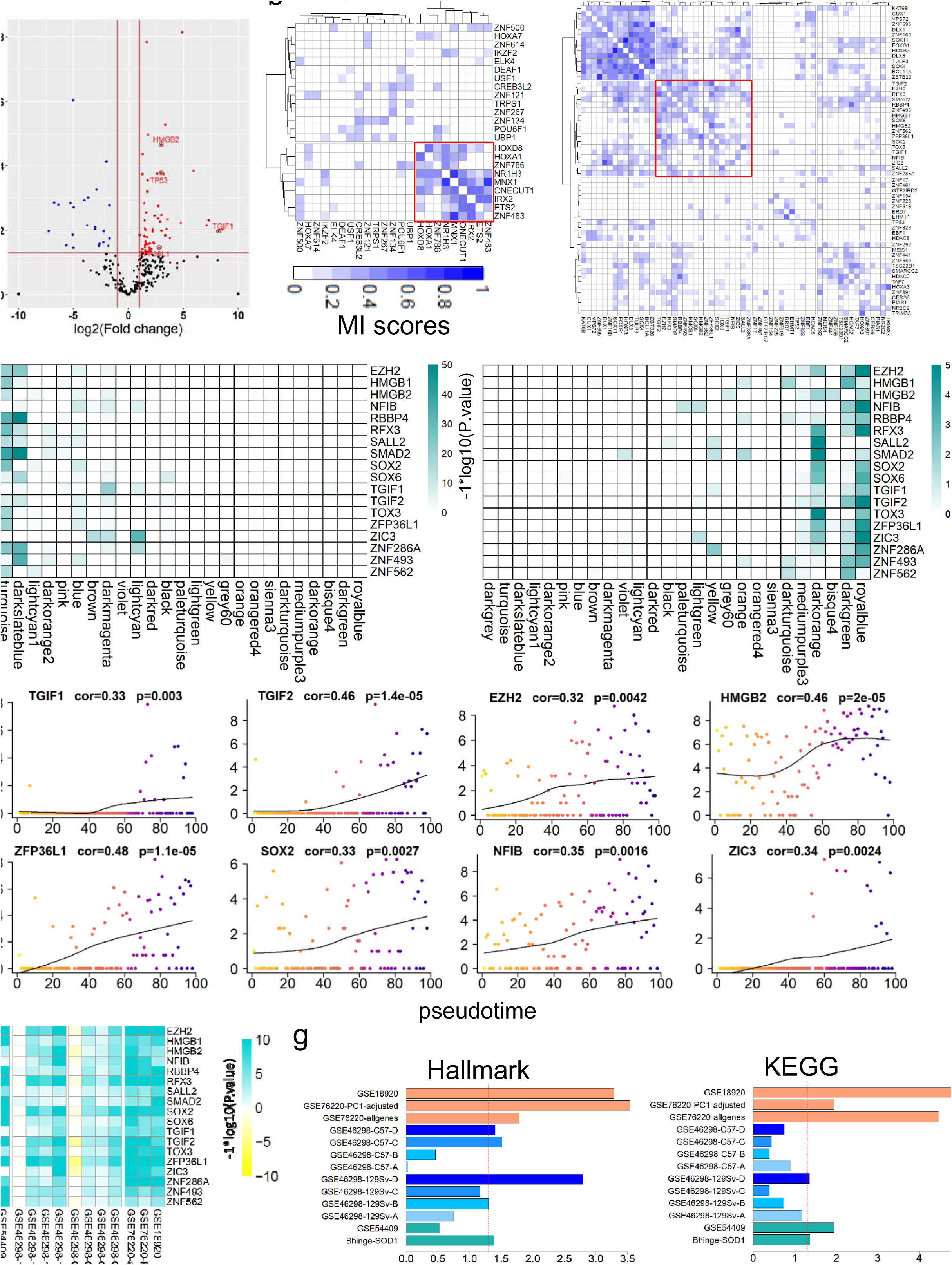
Volcano plots of differentially expressed genes. a) Data displayed for MN. b) Data displayed for IN. Each dot represents a gene. Upregulated genes are coloured red and downregulated genes are coloured blue. Black dots indicate genes not deemed to be significantly differentially regulated. The horizontal red line represents a p-value of 0.01.

**Fig. S4.**
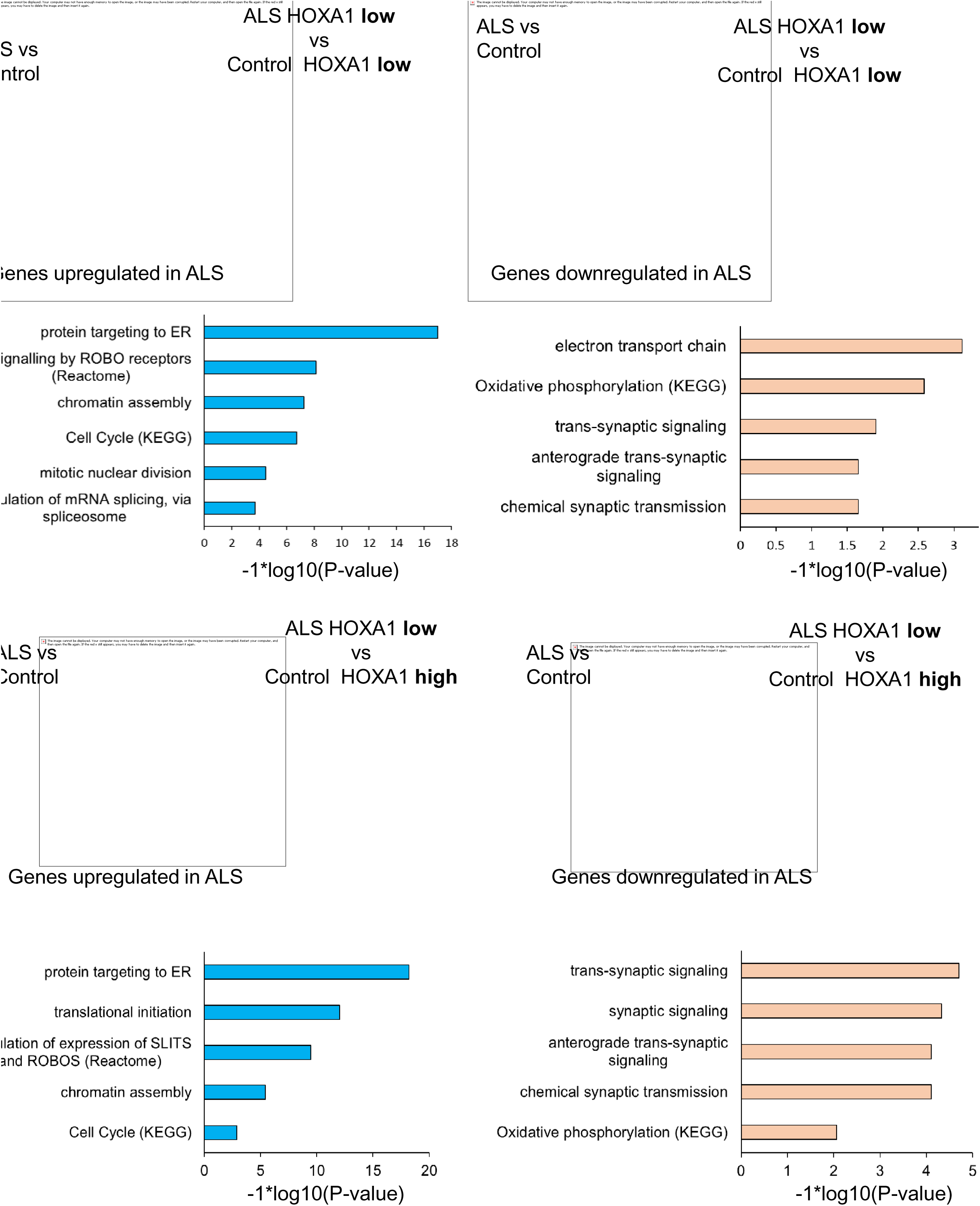
Boxplots displaying module eigengene expression in ALS and Control MN. Module eigengenes (the first principal component of the expression of genes assigned to a module) were used to represent the overall expression of the module. Boxplots show the distribution of the module eigengene values between ALS and Control (CTR) MN for the modules identified to be significantly associated with ALS MN. Positively associated modules: darkgrey, turquoise, darkslateblue, lightcyan1, darkorange2, pink. Negatively associated module: royalblue.

**Fig. S5.**
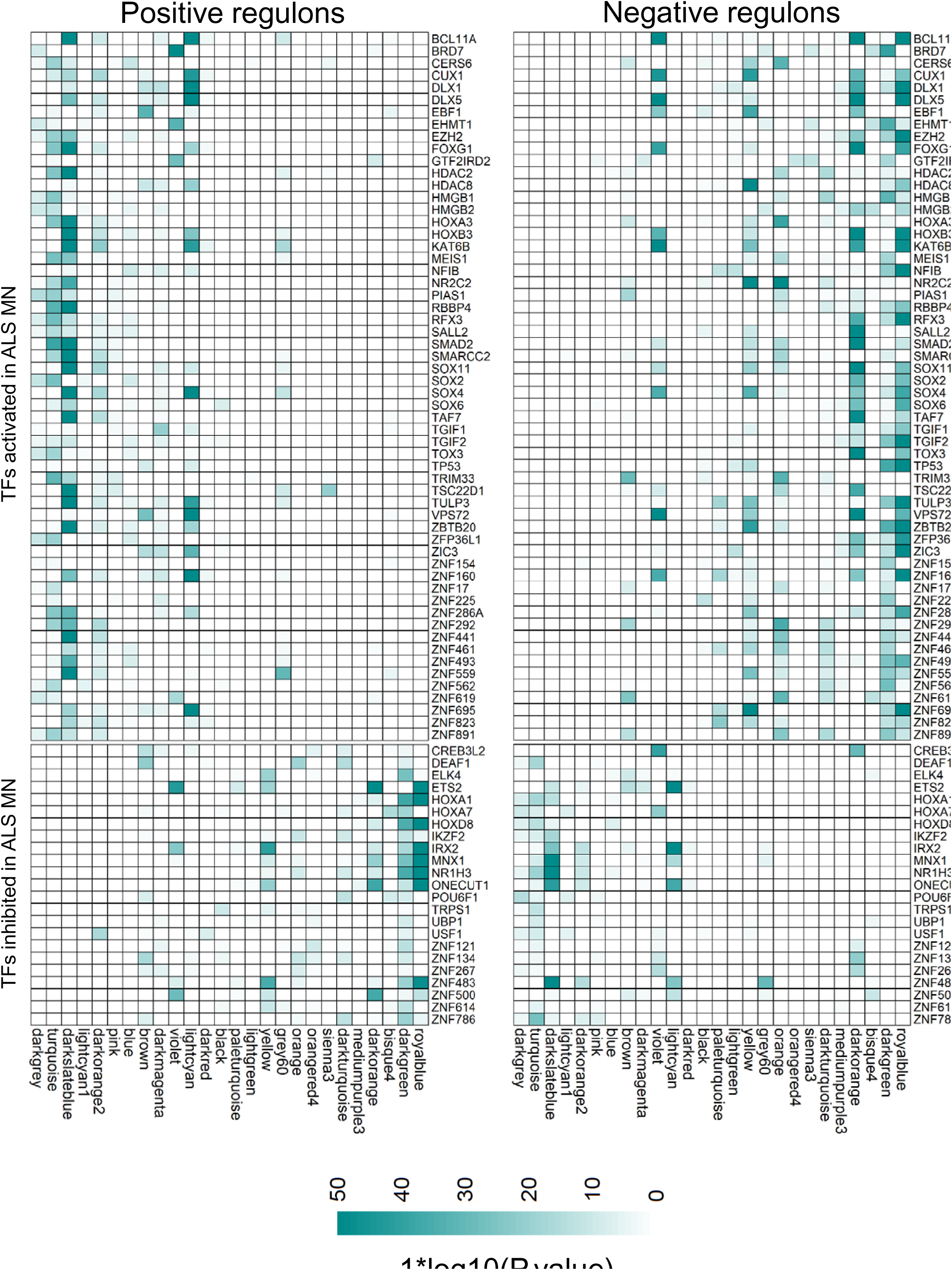
Schematic detailing the master regulator analysis (MRA) a) Generating a transcriptional network using ARACNE. The algorithm estimates pairwise mutual information (MI) scores between TFs and potential target genes, and estimates a significance threshold (p-value) based on a null distribution of scores. MI scores that pass this threshold are deemed valid TF-target pairs. The MI scores do not have a direction i.e. they are always positive. To estimate directionality of regulation, ARACNE estimates correlation between the TF and its target. A positive correlation indicates the TF activates the target while a negative correlation indicates that the TF represses the target. The positive targets of a TF are termed as the positive regulon while the negative targets are deemed the negative regulon. b) Master regulator analysis to identify drivers of ALS MN. Genes found to be differentially expressed in ALS MN are sorted from most upregulated to most downregulated. The algorithm then estimates the enrichment of the positive and negative regulons of each TF across the sorted gene expression dataset. For a given TF, if the positive regulon is enriched in the upregulated end of the sorted genelist and the negative regulon is enriched in the downregulated end, the TF is assigned a positive score. If the inverse is true, the TF is assigned a negative score. TFs with significant positive scores are deemed to be activated in ALS MN and are most likely to drive the ALS MN gene expression. On the other hand, TFs with significant negative scores are deemed to be inhibited in ALS MN. The significance of a score is estimated using random simulations similar to a gene-set enrichment analysis.

**Fig. S6.**
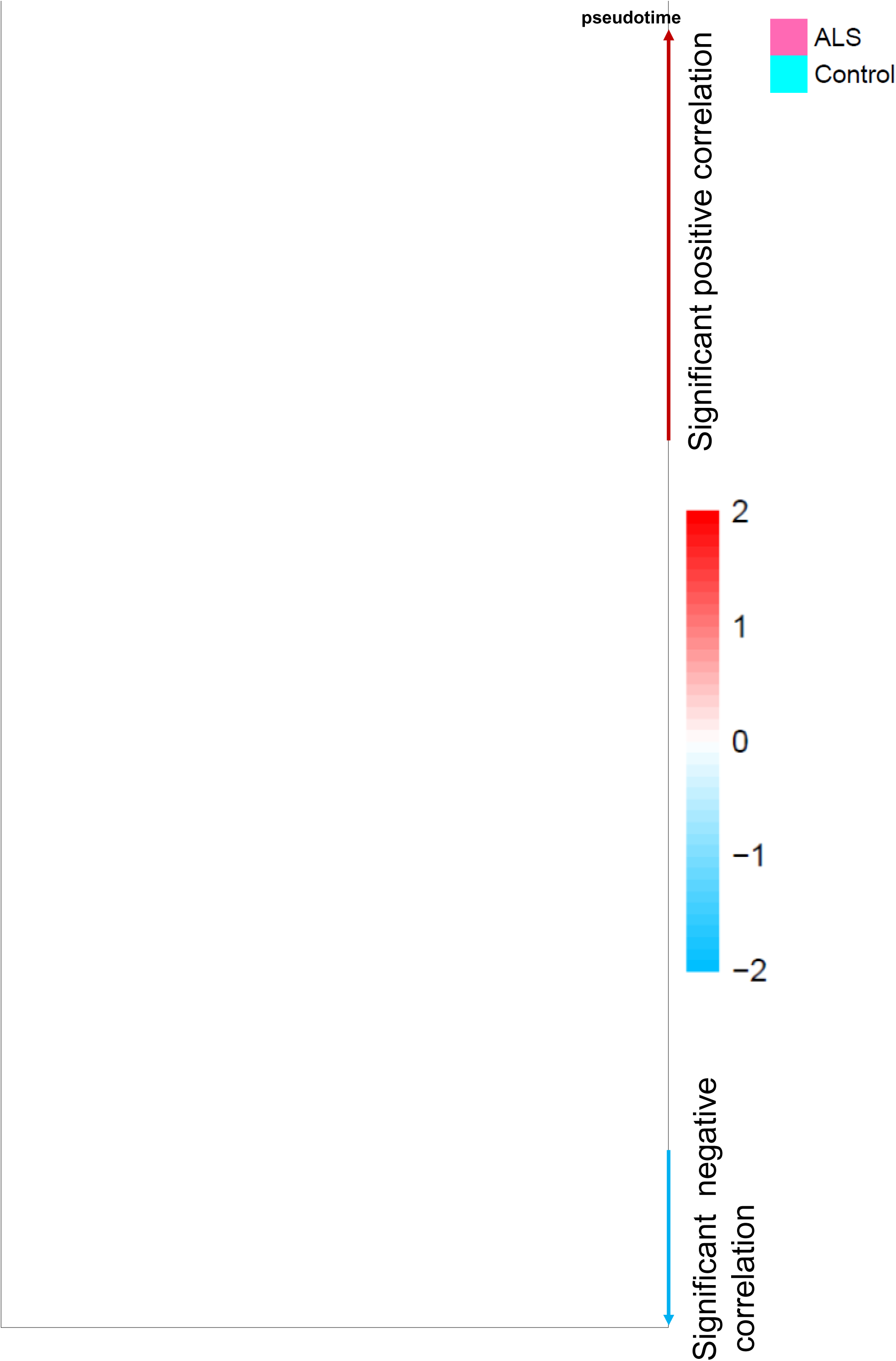
Differential gene expression analysis of MN subgroups based on HOXA1 expression. <u>ALS vs Control</u>: Genes differentially expressed between all ALS MN and all control MN <u>ALS HOXA1 low vs Control HOXA1 low</u>: Genes differentially expressed between ALS MN and Control MN showing low HOXA1 levels. <u>ALS HOXA1 low vs Control HOXA1 high</u>: Genes differentially expressed between ALS MN displaying low HOXA1 levels and Control MN showing high HOXA1 levels. a) Venn diagram showing the overlap between genes upregulated (left panel, p-value = 1e-328) and downregulated (right panel, p-value = 1e-186) between the <u>ALS vs Control</u> group and <u>ALS HOXA1 low vs Control HOXA1 low</u> group. P-values were estimated using a hypergeometric distribution b) GO ontology terms enriched in genes upregulated (left panel, blue bars) and downregulated (right panel, orange bars) in the <u>ALS HOXA1 low vs Control HOXA1 low</u> group. P-values shown were adjusted using the Benjamini-Hochberg procedure. c) Venn diagram showing the overlap between genes upregulated (left panel, p-value = 1e-263) and downregulated (right panel, p-value = 1e-220) between the <u>ALS vs Control</u> group and <u>ALS HOXA1 low vs Control HOXA1 high</u> group. P-values were estimated using a hypergeometric distribution. d) GO ontology terms enriched in genes upregulated (left panel, blue bars) and downregulated (right panel, orange bars) in the <u>ALS HOXA1 low vs Control HOXA1 high</u> group. P-values shown were adjusted using the Benjamini-Hochberg procedure.

**Fig. S7.**
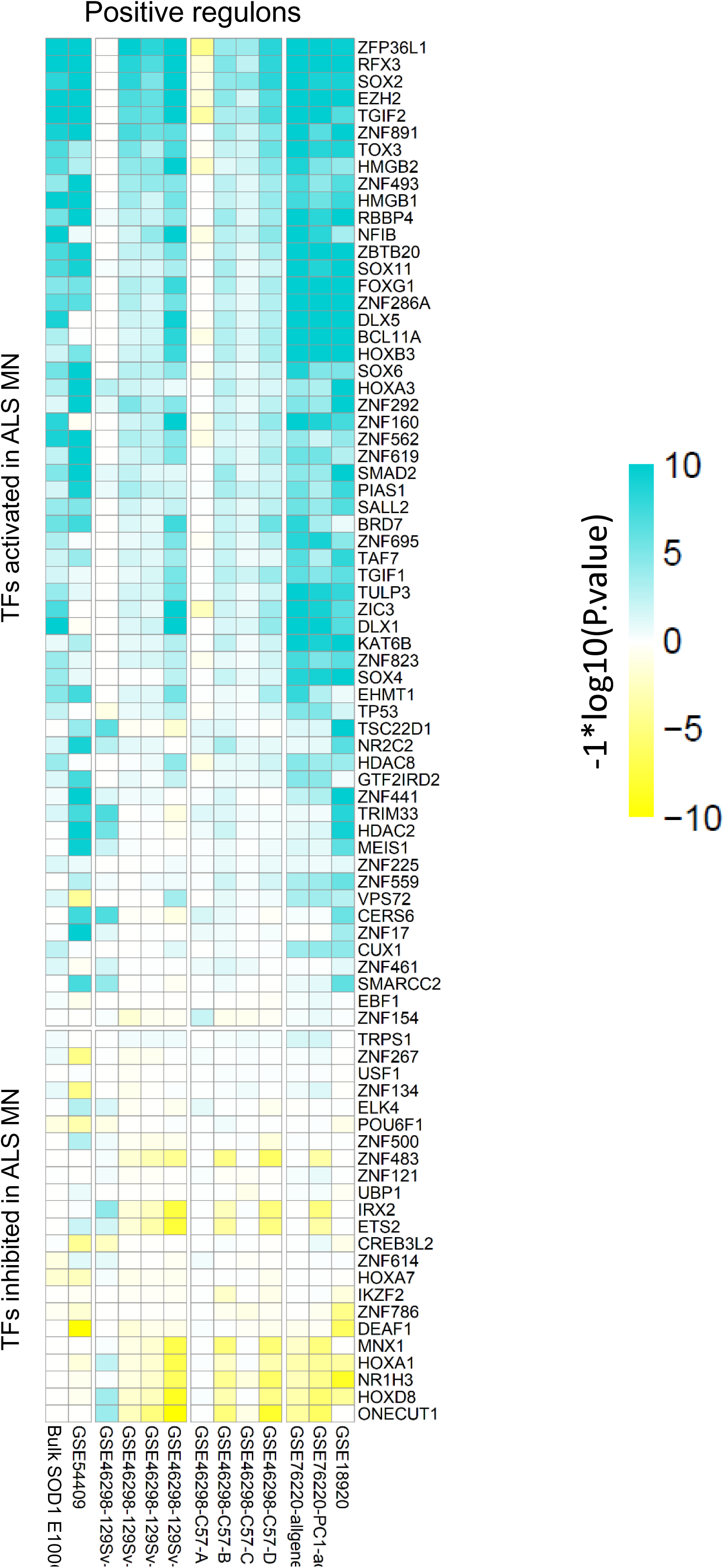
Overlap analysis between regulons of the 81 TFs identified as master regulators of ALS MN gene dysregulation and WGCNA modules. Left panel: analysis using the positive regulons of the TFs. Right panel: analysis using the negative regulons of the TFs. Significance of the overlap was estimated using a hypergeometric distribution. P-values were adjusted using the Benjamini Hochberg procedure, log transformed and plotted as a heatmap.

**Fig. S8.**
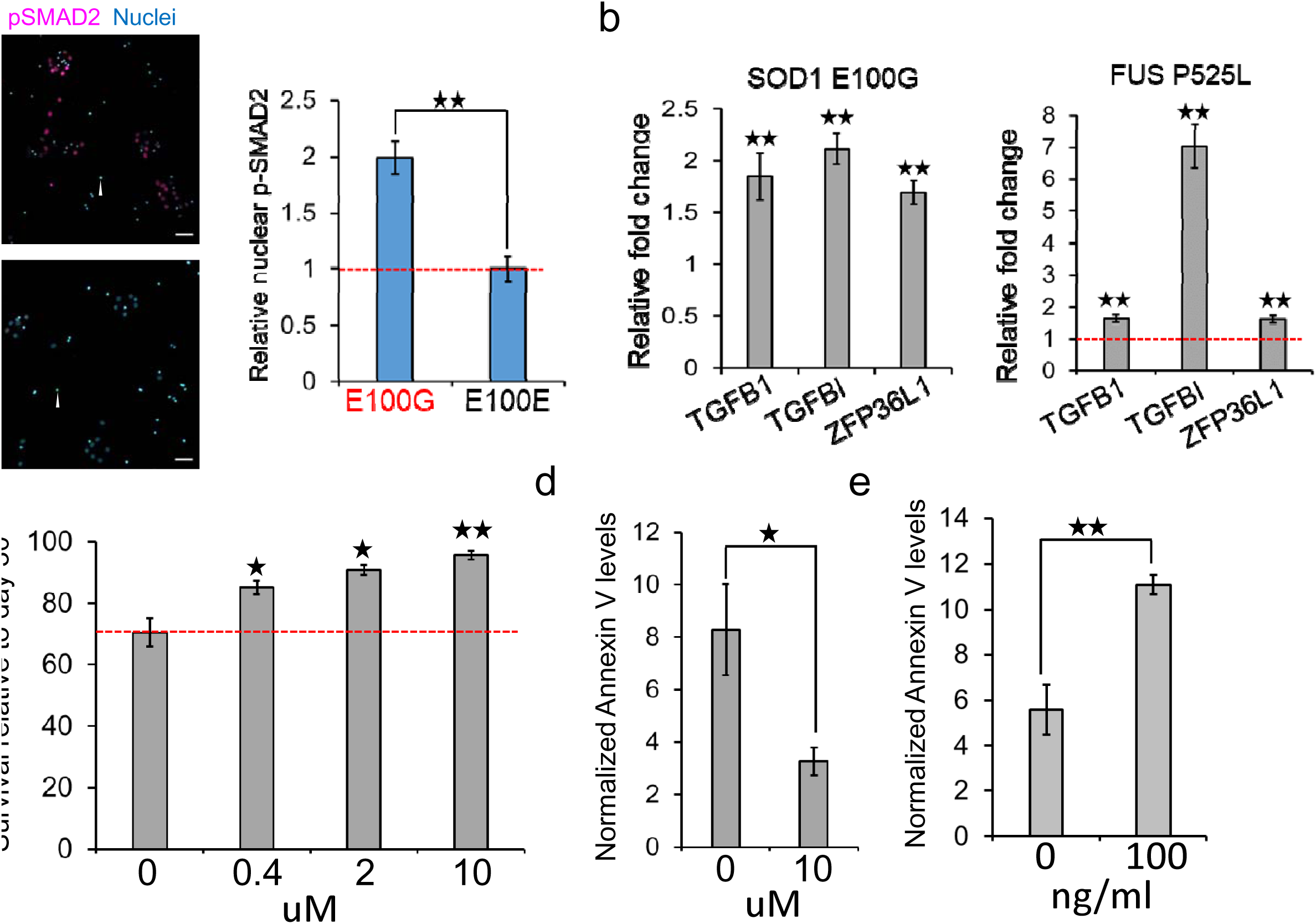
Expression profiles of the 81 master regulator TFs across the MN pseudotime trajectory. Each TF in this set is plotted along the rows. The log normalized values of each TF were mean centered and smoothened across the trajectory. The smoothened expression values were correlated with the pseudotime and sorted. TFs upregulated over the trajectory are at the top while downregulated TFs are at the bottom. X-axis shows the pseudotime. The legend also displays cells that belong to the control (cyan) or ALS (magenta) datasets. TFs that showed significant correlations (adjusted p-value < 0.01) have been highlighted.

**Fig. S9.**
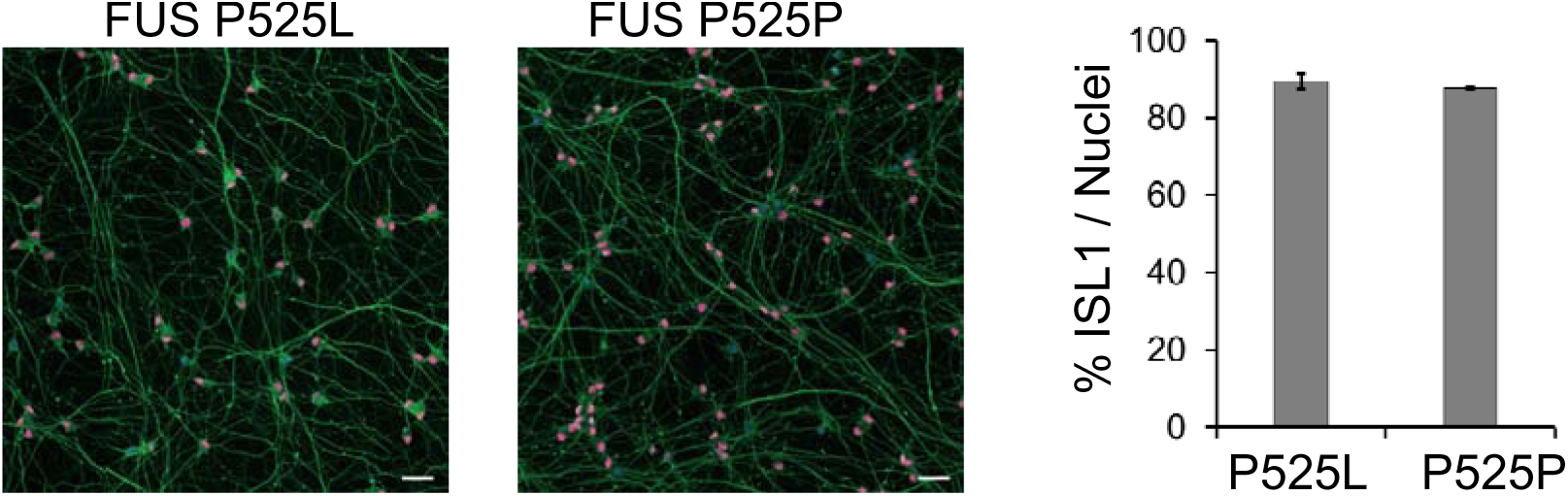
Heatmaps showing the enrichment of the regulons of the 81 master regulator TFs in publicly available ALS datasets. Left panel: Heatmap showing the enrichment of the positive regulons of the master regulator TFs. The datasets are as described in figure 5. Bulk SOD1 E100G: SOD1 E100G iPSC-derived MN analysed in bulk. GSE54409: SOD1 A4V iPSC-derived MN purified via flow sorting. GSE46298-129Sv: MN laser-capture micro-dissected from SOD1 G93A mouse model, fast progressing strain 129Sv. A: pre-symptomatic, B: onset, C: symptomatic, D: end-stage. GSE46298-C57: MN laser-capture micro-dissected from SOD1 G93A mouse model, slow progressing strain C57. A: pre-symptomatic, B: onset, C: symptomatic, D: end-stage. GSE76220: MN laser-capture micro-dissected from sporadic ALS spinal lumbar tissue. GSE76220-PC1: GSE76220 data was filtered for genes involved in wound healing. GSE18920: MN laser-capture micro-dissected from sporadic ALS spinal lumbar tissue. Enrichment of the positive regulons of the TFs were used as input genesets for a GSEA performed on each dataset. Log transformed P-values were assigned the same sign as the GSEA enrichment scores and plotted as a heatmap. Green indicates that the regulon showed a significant positive enrichment while yellow indicates that the regulon showed a significant negative enrichment in the queried dataset of differentially expressed genes. Right panel: same analysis as the left panel using the negative regulons of the TFs.

**Fig. S10.**
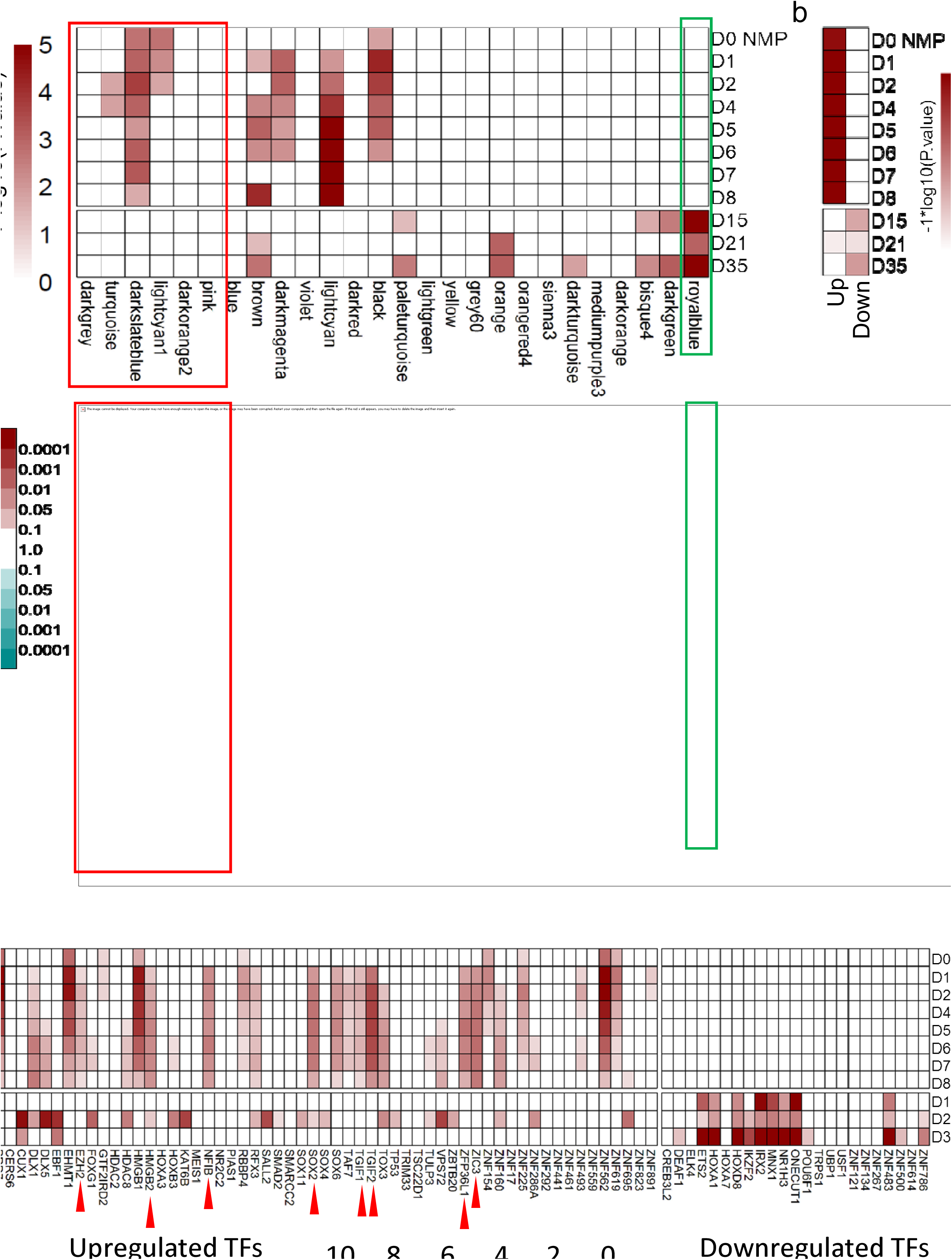
MN differentiated from FUS P525L and isogenic control iPSC. D30 MN were immunostained for the MN marker ISL1 and the pan-neuronal markers TUJ1. Nuclei were stained blue. FUS P525P indicates MN differentiated from the isogenic corrected iPSCs. Scalebar represents 52µm.

**Fig. S11.**
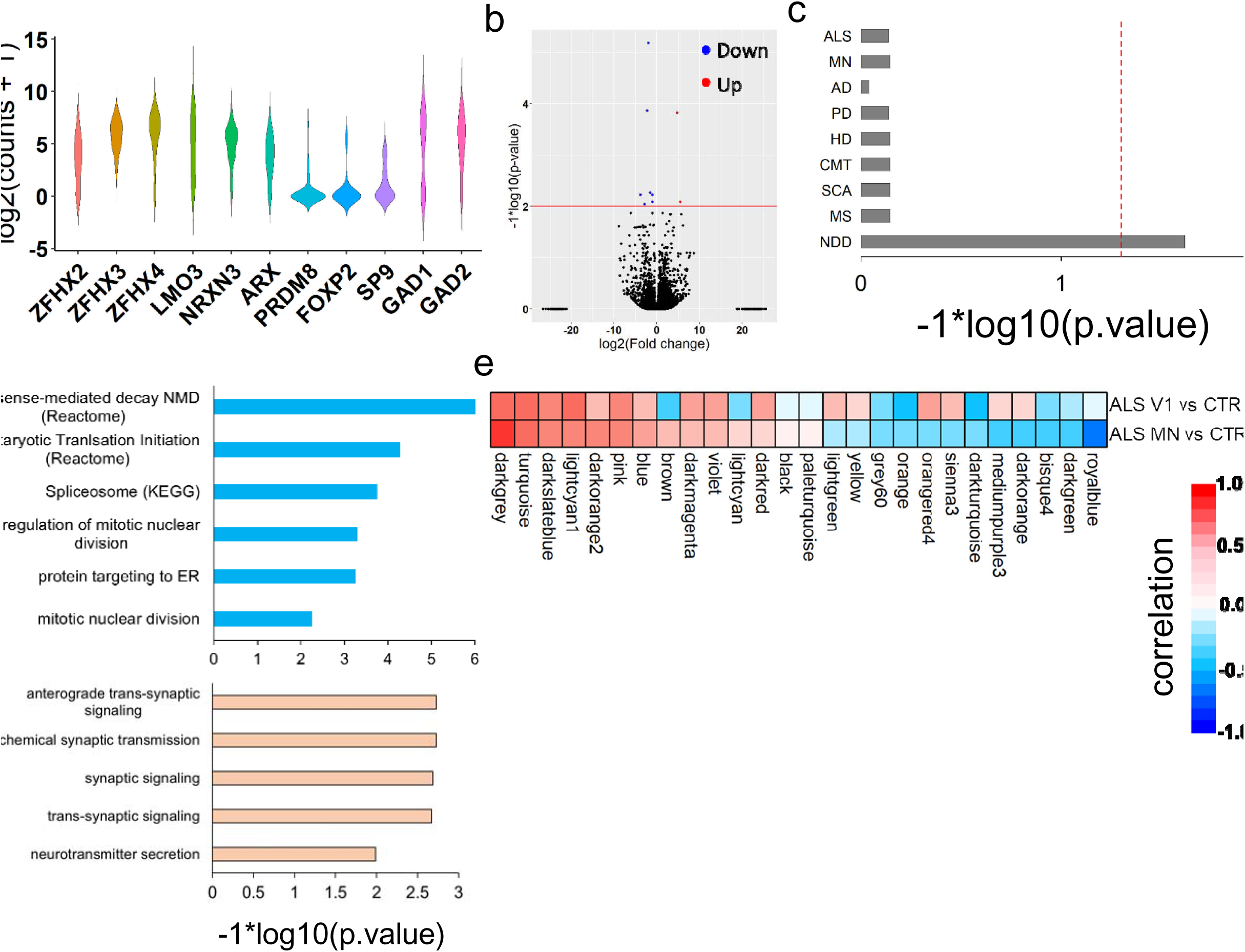
Differential gene expression analysis of ALS V1 IN. a) Violin plot showing log2 transformed expression values of genes known to be expressed in V1 IN. b) Volcano plot of genes differentially expressed in ALS V1 IN. Each dot represents a gene. Upregulated genes are coloured red while downregulated genes are coloured blue. Black indicates genes not deemed to be differentially expressed. Horizontal red line indicates a p-value of 0.01. All p-values were adjusted using the Benjamini Hochberg procedure. c) Enrichment analysis of likely pathogenic variants associated with different diseases from the ClinVar database in genes upregulated in ALS V1 IN. Vertical axis shows terms used to search the ClinVar database to find associated pathogenic variants. ALS: amyotrophic lateral sclerosis, MN: motor neuron, AD: alzheimers disease, PD: parkinsons disease, HD: Huntington’s disease, CMT: charcot marie tooth, SCA: spinocerebellar ataxia, MS: multiple sclerosis, NDD: neurodevelopment disorder. The red dashed line indicates a p-value threshold of 0.05. d) Enrichment of GO terms in genes upregulated (blue bars) or downregulated (orange bars) in ALS V1 IN. The red dashed line indicates a p-value threshold of 0.01. e) Correlation analysis of WGCNA module eigengenes ALS V1 IN and MN. Association of a given module with a specific group (ALS vs control V1 IN or ALS vs control MN) was estimated using Pearson correlations. Red indicates the module is positively correlated with the ALS neurons while blue indicates negative correlation.

**Fig. S12.**
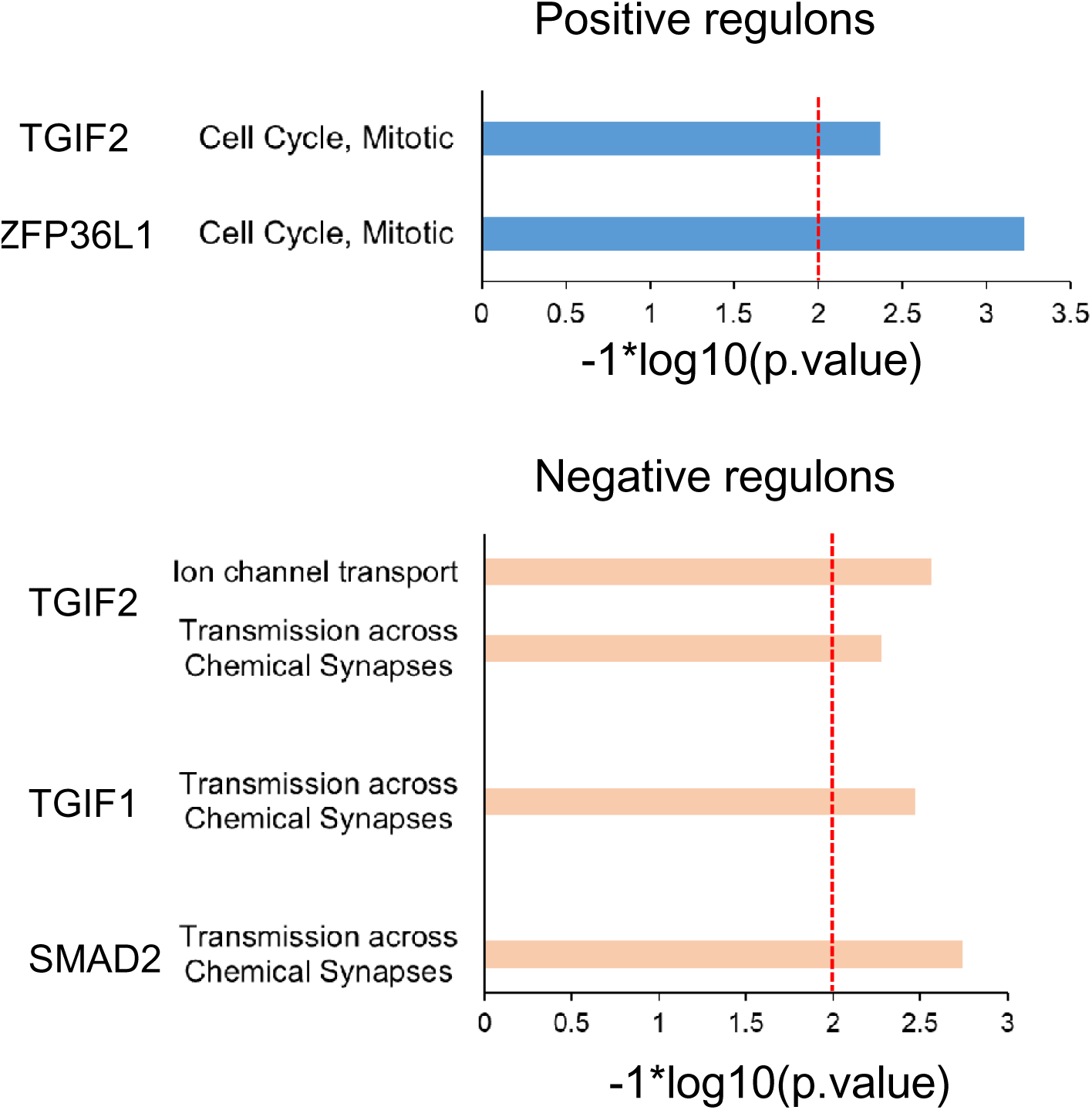
Gene ontology enrichment analysis of the regulons of TGFβ associated TFs.

