## Supplemental-tables for "Single cell transcriptomics identifies master regulators of neurodegeneration in SOD1 ALS motor neurons"

**Table S1. Primary antibodies used for immunofluorescence.**

| **Target** | **Source** | **Catalog number** | **Dilution** |
| --- | --- | --- | --- |
| ISL1 | Abcam | ab8650 | 1:500 |
| ISL1 | Abcam | ab109517 | 1:500 |
| MAP2 | Abcam | ab11267 | 1:1000 |
| NF-H | Sigma | N4142 | 1:1000 |
| Phospho-SMAD2 | Cell signalling technology | 18338 | 1:200 |

**Table S2: Primer sequences used for qPCR analysis.**

| **Gene** | **Forward primer** | **Reverse primer** |
| --- | --- | --- |
| TGFBI | GGACATGCTCACTATCAACGGG | CTGTGGACACATCAGACTCTGC |
| TGFB1 | GTGAGGTCCACGGAAACTGT | TGGCTGGTGCAAAGACATAG |
| ZFP36L1 | CAGGATTCTCTCTCGGACCAG | CAGGCGTCTTGAGTTGTCCA |
| HPRT1 | TGCTCGAGATGTGATGAAGG | AATCCAGCAGGTCAGCAAAG |
| RPL13 | CCTGGAGGAGAAGAGGAAAGAGA | TTGAGGACCTCTGTGTATTTGTCAA |
